# Phosphorylation, disorder, and phase separation govern the behavior of Frequency in the fungal circadian clock

**DOI:** 10.1101/2022.11.03.515097

**Authors:** Daniyal Tariq, Nicole Maurici, Bradley M. Bartholomai, Siddarth Chandrasekaran, Jay C. Dunlap, Alaji Bah, Brian R. Crane

**Affiliations:** Department of Chemistry & Chemical Biology, Cornell University, Ithaca, NY,14853; Department of Biochemistry and Molecular Biology, SUNY Upstate Medical University, Syracuse, NY, 13210; Department of Molecular and Systems Biology, Geisel School of Medicine at Dartmouth, Hanover, NH 03755

## Abstract

Circadian clocks are composed of molecular oscillators that pace rhythms of gene expression to the diurnal cycle. Therein, transcriptional-translational negative feedback loops (TTFLs) generate oscillating levels of transcriptional repressor proteins that regulate their own gene expression. In the filamentous fungus *Neurospora crassa,* the proteins **F**requency (**F**RQ), the **F**RQ-interacting RNA helicase (FRH) and **C**asein-Kinase I (CK1) form the **FFC** complex that represses expression of genes activated by the White-Collar complex (WCC). A key question concerns how FRQ orchestrates molecular interactions at the core of the clock despite containing little predicted tertiary structure. We present the reconstitution and biophysical characterization of FRQ and the FFC in unphosphorylated and highly phosphorylated states. Site-specific spin labeling and pulse-dipolar ESR spectroscopy provides domain-specific structural details on the full-length, 989- residue intrinsically disordered FRQ and the FFC. FRQ contains a compact core that associates and organizes FRH and CK1 to coordinate their roles in WCC repression. FRQ phosphorylation increases conformational flexibility and alters oligomeric state but the changes in structure and dynamics are non-uniform. Full-length FRQ undergoes liquid-liquid phase separation (LLPS) to sequester FRH and CK1 and influence CK1 enzymatic activity. Although FRQ phosphorylation favors LLPS, LLPS feeds back to reduce FRQ phosphorylation by CK1 at higher temperatures. Live imaging of *Neurospora* hyphae reveals FRQ foci characteristic of condensates near the nuclear periphery. Analogous clock repressor proteins in higher organisms share little position-specific sequence identity with FRQ; yet, they contain amino-acid compositions that promote LLPS. Hence, condensate formation may be a conserved feature of eukaryotic circadian clocks.

## Introduction

Eukaryotic circadian rhythms of biological phenomena arise from periodic gene expression patterns that oscillate independent of external cues but can be entrained to light and temperature (Hurley, et al., 2016). At the molecular level, circadian clocks function as transcription-translation negative-feedback loops (TTFLs) that consist of core positive and negative elements (Fig. 1A (Crane, et al., 2014)). In the long-standing model clock of *Neurospora crassa*, the transcriptional repressor complex is known as the FFC, named for its three principal components: Frequency (FRQ, Fig. 1B), the FRQ-interacting RNA helicase (FRH) and Casein Kinase 1 (CK1) (Cha, et al., 2014; Hurley, et al., 2016). FRQ is functionally analogous to Period (PER) in insects and mammals, whereas CK1 is an ortholog of the mammalian CK1 and the Drosophila kinase Doubletime (DBT) (Dunlap, et al., 2017). FRH is found only in the core repressor of the fungal clock. Although FRH is homologous to the yeast helicase MTR4, FRH helicase activity does not appear essential to clock function (Conrad, et al., 2016; Hurley, et al., 2013). The FFC represses the positive arm of the cycle, which involves two zinc-finger transcription factors, White-Collar 1 (WC-1) and White-Collar 2 (WC-2) assembled into the White-Collar Complex (WCC).

**Figure 1:**
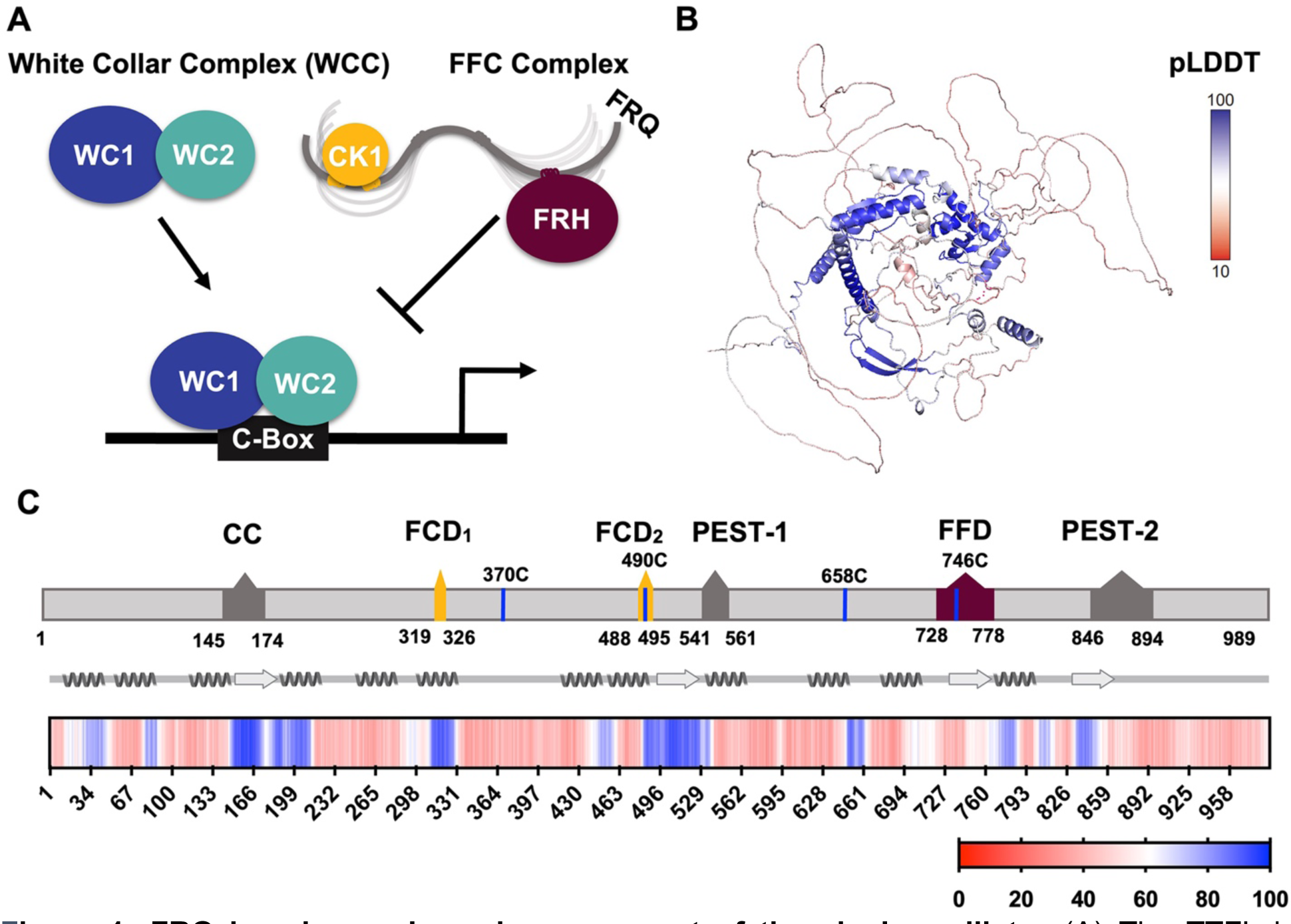
FRQ is a large, dynamic component of the clock oscillator. (A) The TTFL in *Neurospora crassa*. In the core circadian oscillator, **F**RQ associates with **F**RH and **C**K1 to form a repressor complex (the **FFC**) This complex inhibits the positive-acting transcription-factor known as the white-collar complex WCC, composed of White Collar-1 (WC-1) and White Collar-2 (WC- 2). The WCC activates *frq* expression by binding to the c-box region in the *frq* promoter. Stoichiometries of the components are not represented in the schematic. (B) AlphaFold derived model of LFRQ color coded by pLDDT score that highlights the confidence in each predicted region (blue being highest confidence, red lowest). (C) Functional domains of FRQ showing the start position for the LFRQ isoform (AUG_LFRQ_.), key regions and the four native cysteine residues. Predicted secondary structure elements are given below as helices (zigzags) and β-strands (arrows). Color bar shows pLDDT scores across the sequence. Characterized domains include: CC - coiled coil domain, FCD - FRQ:CK1 interacting domain, PEST - proline, glutamic acid, serine, and threonine-rich domain and FFD - FRQ:FRH interacting domain.

WCC binding to the clock-box (c-box) activates *frq* expression (Fig. 1A (Froehlich, et al., 2002)). FRQ production is also highly regulated at the post-transcriptional level through the use of rare codons to retard translation, the expression of an anti-sense transcript and alternative splicing mechanisms (Cha, et al., 2014; Dunlap, et al., 2017; Zhou, et al., 2013). The latter leads to the production of either a long (L-FRQ) or short (S-FRQ) isoform, and the ratio of these isoforms fine tunes period length in response to ambient temperatures (Diernfellner, et al., 2007; Liu, Y., et al., 1997; Zhou, et al., 2013). Translated FRQ binds to FRH via its FFD domain and to CK1 via its two FCD domains (1 and 2) to form the repressive FFC ((Cha, et al., 2014; Dunlap, et al., 2017; Jankowski, et al., 2022); Fig. 1C), although Short Linear Motifs within FRQ may also broadly influence the FRQ-interactome (Pelham, et al., 2021). The FFC enters the nucleus where it directly inhibits the WCC and thereby down-regulates *frq* expression (Hurley, et al., 2013). FRQ- mediated WCC phosphorylation by CK1 (and associated kinases) reduces interactions between the WCC and the c-box to close the feedback loop and restart the cycle (Fig. 1A (Wang, et al., 2019)). FRH facilitates interactions between the FFC and the WCC (Conrad, et al., 2016) and thus, repressor function likely depends upon the FRQ-mediated association of CK1 and FRH. Whereas interaction regions on the FRQ protein sequence have been identified (Fig. 1C), there is little information on the spatial arrangement of the FFC components.

Despite low sequence conservation among the core negative arm proteins across species, they all share a high degree of disorder (Pelham, et al., 2020). FRQ, the analogous Drosophila (d) Period (PER) protein and the analogous mammalian (m) PER have > 65% disorder predicted in their sequences with ∼86% of the FRQ sequence assumed to have no defined structure (Pelham, et al., 2020). Both FRQ and mPER2 have biochemical properties characteristic of intrinsically disordered proteins (IDPs) (Hurley, et al., 2013; Marzoll, et al., 2022). These clock repressors also undergo extensive posttranslational modification (PTM), particularly phosphorylation (Dunlap, et al., 2018; Pelham, et al., 2020). FRQ acquires over 100 phosphorylation sites over the course of the circadian day (similar to PER phosphorylation by DBT), predominantly due to CK1 (Baker, et al., 2009; Tang, et al., 2009). The phosphorylation events have two consequences: 1) those that weaken the interaction between FRQ and the WCC or CK1 (Cha, et al., 2014; Liu, X., et al., 2019); and 2) those that direct FRQ down the ubiquitin-proteasome pathway (Larrondo, et al., 2015), although only the former are required for circadian rhythmicity (Larrondo, et al., 2015). Multi-site phosphorylation of FRQ influences its inhibitory function and clock period length (Larrondo, et al., 2015; Liu, X., et al., 2019). Although the consequences of phosphorylation-driven conformational changes in FRQ have been probed by proteolytic sensitivity, the challenges of producing FRQ have limited direct assessment of its conformational properties (Liu, X., et al., 2019; Pelham, et al., 2020; Querfurth, et al., 2011). IDPs such as FRQ are known to participate in liquid-liquid phase separation (LLPS), which has emerged as a key mechanism for organizing cellular components, coordinating metabolism and directing signaling through, for example, the formation of membraneless organelles (MLOs) (Dignon, et al., 2020; Wright, et al., 2014). Emerging studies in plants (Jung, et al., 2020), flies (Xiao, et al., 2021) and cyanobacteria (Cohen, et al., 2014; Pattanayak, et al., 2020) implicate LLPS in circadian clocks, and in *Neurospora* it has recently been shown that the Period-2 (PRD-2) RNA-binding protein influences *frq* mRNA localization through a mechanism potentially mediated by LLPS (Bartholomai, et al., 2022). Intriguing questions surround why clock repressor proteins are so disordered, undergo extensive post-translational modification and whether LLPS has a role to play in their function.

Clock proteins and their complexes have been challenging to characterize due to their large size (>100 kDa), low sequence complexity, extensive disorder and modification states. Herein, we reconstitute FRQ and the FFC to assess the global and site-specific structural properties of these moieties, as well as the dependence of their properties on multi-site phosphorylation. FRQ undergoes non-uniform conformational change upon phosphorylation and arranges FRH close to CK1 in the FFC. We also demonstrate that FRQ indeed participates in phosphorylation-dependent LLPS in vitro, and as a condensate sequesters FRH and CK1 while altering CK1 activity. Our results suggest how a massively disordered protein such as FRQ organizes the clock core oscillator and reveal the effects of extensive phosphorylation on the conformational and phase-separation properties of a large IDP.

## Results

### Sequence assessment and general properties of FRQ

Analysis of an AlphaFold 2.0 (Jumper, et al., 2021) model of full-length FRQ indicates that FRQ lacks a defined arrangement of large folded domains; instead, most of the molecule is unstructured (Fig. 1B), in agreement with previous studies predicting that FRQ mostly comprises intrinsically disordered regions (Hurley, et al., 2013). However, FRQ is also predicted to contain a small, clustered core, with several secondary structural elements that collapse together to form a compact nucleus surrounded by long flexible disordered regions (Fig. 1B). The AlphaFold pLDDT scores, which represent the confidence in the structural prediction (Jumper, et al., 2021), are very low across the model, except within the core (Fig. 1B, Fig. S1), which contains the CK1 binding region. The FRH binding region, FFD, falls within a more flexible region with a lower pLDDT score than that of CK1. Helices compose the majority of secondary structural elements predicted by AlphaFold (Fig. 1B). The assignments of the helical regions were well supported by the Agadir program that predicts helical propensity based on the helix/coil transition theory (Muñoz, et al., 1994). Four regions distributed throughout the length of FRQ (residues 194-212, 312-329, 709-719 and 801-817) have at least 10% helical propensity and these helical regions mostly agree with those of the AlphaFold model (Fig. 1B). Together, these analyses primarily indicate that FRQ assumes a broad conformational ensemble containing an ordered nucleus that interacts extensively with partially ordered and flexible regions. However, much of the structural details of the model, including the positioning and interactions between secondary structural elements, are predicted with low confidence, thereby underscoring the need for a multi-pronged biophysical study of the protein.

### Reconstitution of FRQ in unphosphorylated and highly phosphorylated states

The size and intrinsic disorder of FRQ have made it challenging to recombinantly express and purify, thereby limiting investigations of its biophysical properties. Previous purifications have characterized FRQ as an IDP, but suffered from low yields and aggregated species (Hurley, et al., 2013; Lauinger, et al., 2014). We also encountered extensive protein degradation and aggregation with conventional purification methods and affinity tagging strategies; however, methods proven effective for IDPs (Pedersen, et al., 2020) yielded comparatively well-behaved protein. Briefly, we carried out nickel affinity chromatography and size exclusion chromatography (SEC) under denaturing conditions to maximize yield and reduce aggregation of full-length L-FRQ (residues 1-989; Fig. 2A) (Seal, et al., 2021). Solubilization with high concentrations of guanidinium-HCl prevented proteolysis, aggregation and limited co-purification of impurities that were otherwise difficult to remove. Once FRQ was isolated, the protein remained soluble, monodisperse and could be concentrated after the denaturant was removed by dialysis (Fig. 2A,B). To our knowledge, FRQ is one of the largest native IDPs produced in amounts amenable to structural and biophysical characterization.

**Figure 2:**
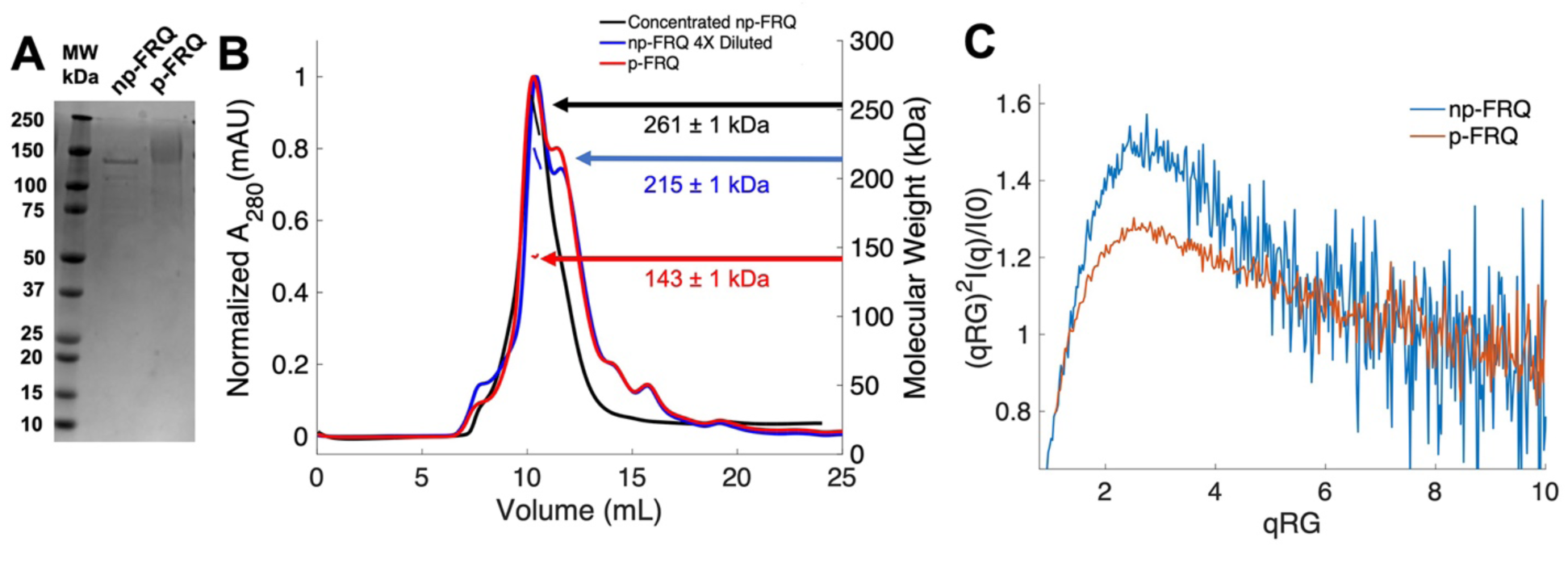
Purification of FRQ in non-phosphorylated and phosphorylated forms reveals IDP behavior. (A) SDS-PAGE gel showing non-phosphorylated FRQ (np-FRQ) and phosphorylated FRQ (p-FRQ) alongside a molecular weight ladder. Both samples represent the purified full-length long isoform (L-FRQ, residues 1-989) in non-denaturing buffer. (B) Size exclusion chromatography–multiangle light scattering (SEC-MALS) of phosphorylated FRQ (p-FRQ) and non-phosphorylated (np-FRQ). The molecular weight (MW) of p-FRQ (∼143 ± 1 kDa) is somewhat larger than that of an unphosphorylated monomer (∼120 kDa), but substantially smaller than that of a dimer, whereas the high concentration MW value of np- FRQ (∼261 ± 1 kDa) is slightly larger than that expected for a dimer (∼240 kDa). Note that the p-FRQ MW is affected by >80 phospho sites. The shoulder to the right of the main peak had low light-scattering and a large MW error. The colored arrows show the MW of each sample as determined by the MALS traces, which are inlaid within the SEC peaks. (C) Dimensionless Kratky plot of SAXS data for p-FRQ and np-FRQ, in both cases the peak positions of the curves are shifted from that expected for a globular protein (√3,1.104), but neither plateau at 2, which would be characteristic of a flexible, denatured polypeptide (Durand, et al., 2010). See Table S1 for additional SAXS parameters. Total protein concentration was between 50-75 μM for the MALS and SAXS experiments.

To generate phosphorylated FRQ, the protein was co-expressed with (untagged) CK1, which forms a stable complex with FRQ and acts as its primary kinase (Querfurth, et al., 2011). The same denaturing purification procedure was followed as for FRQ expressed without CK1. Phosphorylation was confirmed visually by altered migration on SDS-PAGE (Fig. 2A) and mass spectrometry (Fig. S2). The latter revealed a network of > 80 phosphorylation sites (Fig. S2), most of which agree with those found on natively expressed FRQ (Baker, et al., 2009; Tang, et al., 2009). Hereafter, we refer to this preparation as phosphorylated FRQ (p-FRQ) whereas non-phosphorylated FRQ (np-FRQ) refers to FRQ expressed without CK1. In general, we found that co-expression of FRQ with CK1 increased protein yield and stability, while limiting aggregation and degradation.

### Global properties of FRQ

Size exclusion chromatography coupled to multi angle light scattering (SEC-MALS) of p-FRQ gives a molecular weight measurement consistent with a primarily monomeric state (MW of ∼143 kDa versus a predicted MW of ∼110 kDa; Fig. 2B). In contrast, the np-FRQ measured by MALS indicates a dimer and its measured mass distribution increases upon concentration (Fig. 2B) Hence, purified np-FRQ dimerizes, whereas phosphorylation destabilizes this association. SEC-coupled small-angle x-ray scattering (SAXS) analyses revealed that both p-FRQ and np-FRQ shared characteristics of highly flexible proteins, but that p-FRQ was more extended than np-FRQ (Table S1). The dimensionless Kratky plots of p-FRQ and np-FRQ, which report on compactness and conformational dispersity (Durand, et al., 2010; Martin, Erik W., et al., 2021; Sagar, et al., 2020; Trewhella, et al., 2017), revealed high degrees of flexibility for both proteins (Fig. 2C). The peak values of the plot were shifted from those expected for compact structures but were also not indicative of fully disordered proteins (Durand, et al., 2010; Rambo, et al., 2011; Sagar, et al., 2020). Furthermore, p-FRQ appears more disordered than np-FRQ, a characteristic also reflected in their pairwise distance distributions (Fig. S3). These data then indicate that FRQ generally adopts a relatively monodisperse conformational state that is partially compact but highly flexible and then increases expanse and flexibility upon phosphorylation. This general picture is consistent with the increased susceptibility to proteolysis that FRQ experiences with phosphorylation (Liu, X., et al., 2019; Pelham, et al., 2020; Querfurth, et al., 2011).

### Local measures of FRQ structure and dynamics

The properties of FRQ that include its intrinsic disorder and propensity to form heterogenous ensembles are not well suited to commonly applied structural characterization methods. We applied site-specific spin labeling (Fig. 3A) and electron spin resonance spectroscopy (ESR) to monitor FRQ local structural properties. FRQ has four native cysteine residues that provide reactive sites for nitroxide spin-labels (Fig. 3A,B (Berliner, et al., 1982)). The Cys residues distribute relatively evenly throughout the protein sequence and fall within areas of varying structural order that include the binding regions for the core clock proteins (FRH, CK1, WC1 and WC2; Table S2-3; Fig. 3B). Furthermore, we developed and applied enzymatic SortaseA and AEP1 peptide-coupling methods (Chandrasekaran, et al., 2021; Nguyen, et al., 2015) to spin-label FRQ at its N- and C- termini (Fig. 3A; Fig. S4).

**Figure 3:**
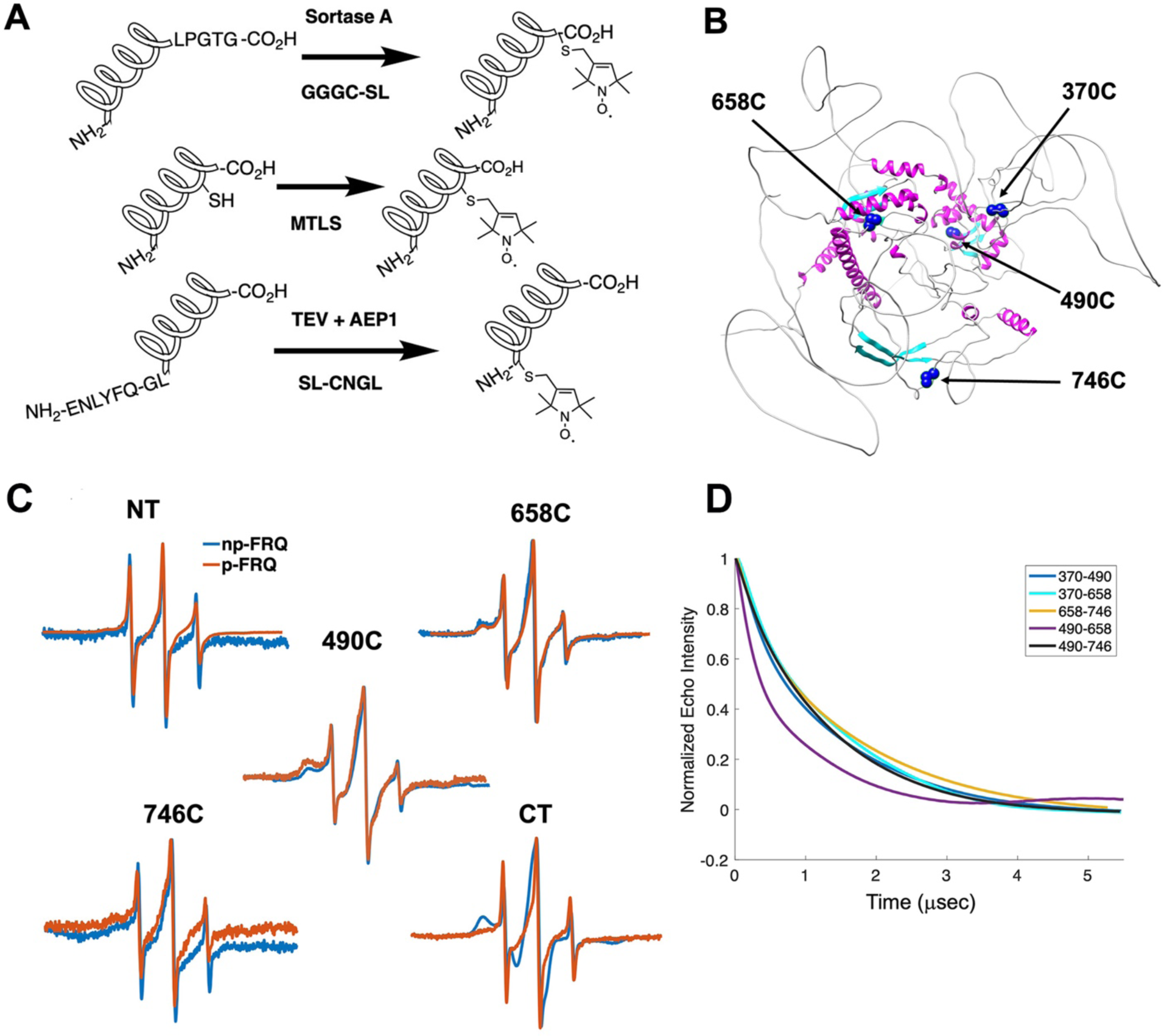
**ESR spectroscopy studies of spin-labeled FRQ**. (A) Spin-labeling strategies for FRQ. MTSL was used for internal labeling of cysteine residues and AEP1 and SortaseA were used to label the N and C terminus, respectively. TEV protease cleavage was used to reveal the AEP1 ligation site. (B) AlphaFold generated model of FRQ with the positions of native spin-labeled cysteine residues highlighted in dark blue. (C) X-band (9.8 GHz) room temperature cw-ESR measurements for various spin-labeled FRQ variants, each containing a single Cys labeling site, as designated. Data represented as first-derivative spectra with the horizontal axes depicting the static field strength (typically 3330 to 3400 G), and the vertical axis depicting the change in magnetic susceptibility with respect to the static field, in arbitrary units. (D) Time-domain data of DEER measurements for double-cysteine labeled FRQ variants. The baseline-corrected time domain dipolar evolution trace fits are shown; raw data and baseline corrections are given in Fig. S5-6. The 490-658 separation is noticeably closer and more ordered than the others. The total protein concentration was between 50-75 μM for all experiments.

Line-shape analysis from continuous wave (cw)-ESR spectroscopy reports on local dynamics (Franck, et al., 2015; Freed, 2005). For rapidly rotating nitroxide spin labels (SLs), the characteristic spectrum consists of three narrow lines of equal width and intensity. As the rotational mobility decreases, the lines broaden, especially the high-field line (Bonucci, et al., 2020; Emmanouilidis, et al., 2021). Cw-ESR reveals that FRQ has regions of varying order that agree with computational predictions and that some of these sites undergo phosphorylation-dependent conformational changes (Fig. 3C). Spin-labels attached to Cys490 and Cys658 (490SL and 658SL) are predicted to be within an α-helix and a β-strand respectively. These sites correspondingly showed a broad low-field (left-most line) component and a small high-field component (right-most line) indicative of slower tumbling compared to sites predicted to be in flexible regions, such as 746SL (Fig. 3C). The 490SL site had a broader profile compared to that of the 658SL site, although both are relatively well ordered. We observed a similar trend with δ, which is the peak-to-peak width of the central line (Hubbell, et al., 2000). 490SL and 658SL had similar δ values that were higher than that of 746SL, indicating that they are less flexible than 746SL (Table S4). Furthermore, the spectra also highlighted structural differences between the phospho forms: p-FRQ and np-FRQ gave very different δ values at 490SL and the C-terminus, thereby demonstrating how phosphorylation can influence conformational properties site-specifically. The C-terminus may have become more dynamic in p-FRQ as compared to np-FRQ, because the C-terminus contains a cluster of phospho-sites (Fig. 3C). In contrast, the N-terminus and especially 490SL, which locates to the CK1-interaction region of the FCD_2_ domain, are more ordered in p-FRQ. Such differences in mobility between p-FRQ and np-FRQ in the FCD (490SL) may reflect changes in FRQ:CK1 interactions as FRQ becomes progressively phosphorylated across the circadian period (Liu, X., et al., 2019).

Double electron–electron resonance (DEER) spectroscopy provides distance distribution between two nitroxide labels in the range of 15 Å to ∼100 Å. We produced 6 cysteine pair variants that each contained only two of the four native FRQ Cys residues to measure pair-wise distances and one single Cys variant to test for oligomerization. Owing to association and aggregation when concentrated, np-FRQ was not amenable to DEER measurements; however, such issues were not prohibitive with p-FRQ. Nevertheless, due to the challenges of spin-labeling appreciable concentrations of p-FRQ, the modulation depths of the DEER data, which reflect the number of spins in range to produce specific dipolar interactions, are small (∼2-4%). These values compare to ∼10-12% for a well-ordered, highly labeled pair of sites under these instrument conditions (Dunleavy, et al., 2023). The single Cys spin-labeled variant 490SL produced log-linear time domain traces indicative of a uniform background with little evidence of dimerization, consistent with the MALS data showing p-FRQ to be monomeric (Fig. S5). A second single Cys variant, 746C likewise produced little dipolar signal, although for this variant, solubility issues limited spin concentration (*Fig. S5*). The double 370SL:746SL p-FRQ variant also gave no dipolar signals and hence no dimerization as reported by those two sites (Fig. 3D). When at least one of the two spin labels resides outside of the predicted core, the DEER experiments of the spin pairs generally revealed small, albeit significantly slow decaying time domain traces that deviate from background (Fig. 3D; Fig. S5-6). Such traces are indicative of long inter-spin distances and conformational flexibility, which would be consistent with the large mobility and intrinsic disorder of these peripheral regions (Fig. S5-6). Only variant 490SL:658SL gave a fast-decaying time-domain signal, indicative of short inter-spin distances and conformational rigidity (Fig. S5-6). Despite the overall uncertainty in the AlphaFold model (Fig. 1B), it is notable that 490SL and 658SL both fall within the predicted ordered core.

### FFC reconstitution and component arrangement

The FFC was generated by mixing p-FRQ or np-FRQ with excess FRH and CK1, followed by SEC purification to give p-FFC or np-FFC, respectively. FRHtiN (residues 100-1106), which binds FRQ, was used to improve expression and enhance stability (Conrad, et al., 2016; Hurley, et al., 2013). The resulting symmetric SEC traces and SDS-PAGE gels revealed pure and homogenous ternary complexes in each case (Fig. 4A). As expected, CK1 phosphorylates FRQ in np-FFC (Fig. 4A, Fig. S7), and this activity does not require priming by other kinases (Marzoll, et al., 2022). SEC-SAXS analysis of the FFC showed that both FRQ phospho-forms produced complexes that were more globular than FRQ alone, but still somewhat flexible, with the p-FFC being more flexible than the np-FFC (Fig. 4B, Fig. S3). MW estimates of the p-FFC and np-FFC from SAXS indicate a 1:1 stoichiometry of FRQ:FRH, with a likely single component of CK1 also contained within the complex (Table S1).

**Figure 4:**
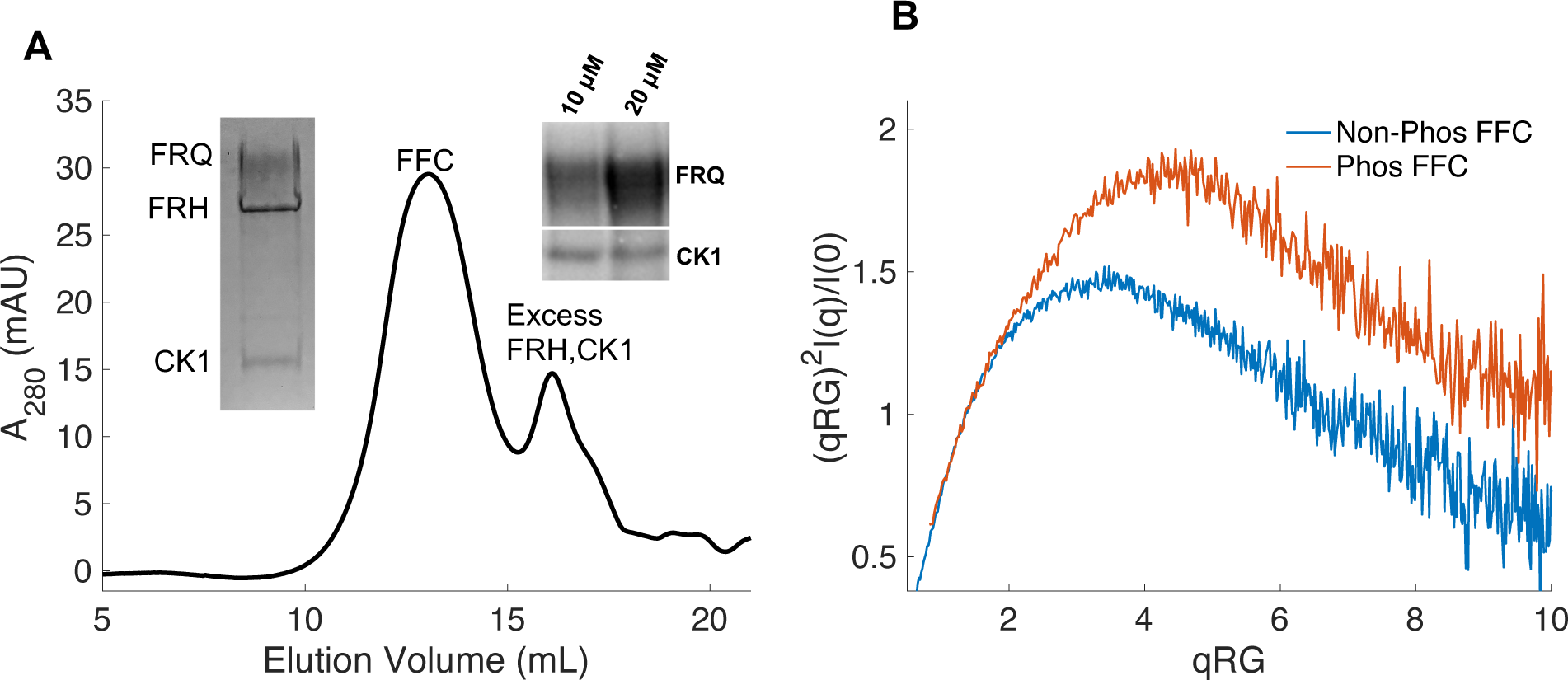
**Properties of the FFC**. (A) SEC of the FFC; left inset: SDS-PAGE gel inset of the components; right inset: ^32^P-autoradiogram of FRQ phosphorylation by CK1 and CK1 autophosphorylation at 10 and 20 μM FRQ all in the presence of FRH. Complete autoradiograms are shown in Fig. S7. (B) Overlay of dimensionless Kratky plots of SAXS data from the FFC formed with either np- or p-FRQ (protein concentrations were between 50-75 μM). The SAXS-derived MW of np-FRQ FFC = 285 kDa, which compares to a predicted value of 273 kDa; MW of p-FFC = 340 kDa; Table S1).

DEER measurements from selectively spin-labeled components within the complex were used to probe its overall architecture (Fig. 5; Fig. S8-S11). An ADP-βSL probe (Muok, et al., 2018) bound to both FRH and CK1 and revealed that the FRH and CK1 ATP-binding pockets are closely positioned within the FFC complex (Fig. 5A; Fig. S8). Nonetheless, the presence of broad and multiple peaks in the distance distributions indicated conformational heterogeneity. Spin labels placed within the FCD2 and FFD domains of FRQ (490C and 746C), which bind CK1 and FRH respectively, produced a distance distribution similar to that of the labels that target the ATP binding sites of the two enzymes (Fig. 5A) (Cheng, et al., 2005). These independent observations support the conclusion that FRQ organizes FRH and CK1 in close proximity within the FFC. Notably, purified FRH and CK1 do not interact in the absence of FRQ (Fig. S8B).

**Figure 5:**
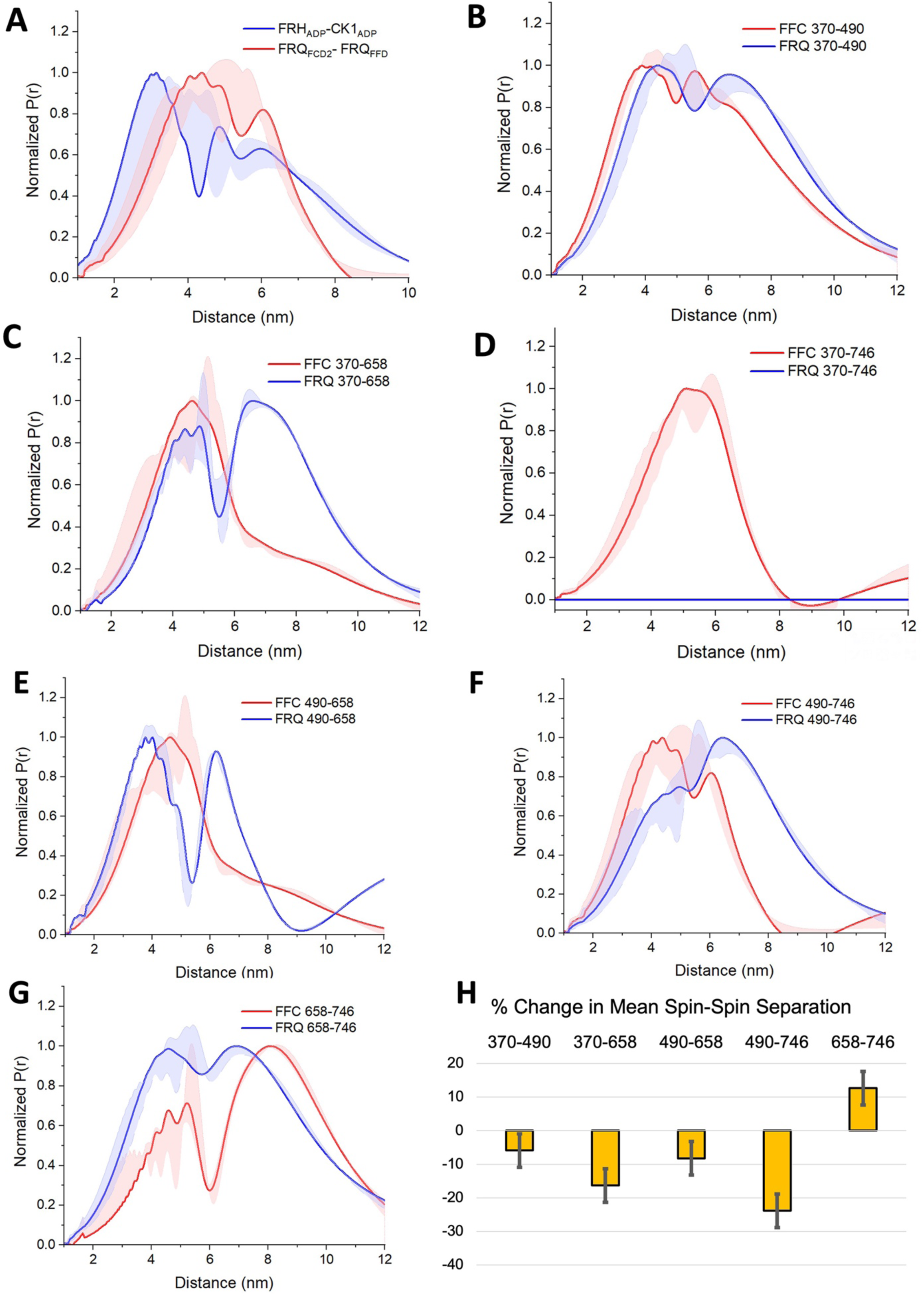
PDS distance distributions of spin-labeled FRQ and the FFC. (A) DEER-derived distance distributions of the FFC wherein the spins were directed by ADP linkage to target the FRH and CK1 ATP-binding sites (blue) or in the form of MTSL labeling of cysteine residues within the FCD2 and FFD domains of FRQ (red). Uncertainties in the distributions at each distance point (see Supplementary Materials and Methods) are represented by the widths of the blue and red shading. Time-domain data and distance distributions with error estimates for single samples are shown in Fig. S8. (B-G) DEER-derived distance distributions of cysteine residue pairs labeled with MTSL within p-FRQ, either alone (blue) or within the FFC (red). Uncertainties in the distributions at each distance point are represented by the widths of the blue and red shading. Time-domain data and distance distributions with error estimates for single samples are shown in Fig. S5, S9-S10. (H) Percentage change in the average separation of FRQ when it binds FRH and CK1. Errors are derived from the uncertainties in the distance distributions (also shown in Fig. S9-10). In these experiments, only FRQ was labeled with MTSL at the positions noted. Note: Total protein concentrations were between 50-75 μM for all experiments.

Spin-label separations within p-FRQ changed substantially when FRH and CK1 bound (Fig. 5; Fig. S5-6,S9-11). In general, the time domain decays were more rapid in the FFC compared to p- FRQ alone and the resulting distance distributions had smaller mean distances and less breadth (Fig. 5B-H). However, the changes were not uniform; for example, the 658SL-to-746SL distance distribution increased in the complex compared to free FRQ (Fig. 5G). To further assess how the distributions of p-FRQ conformations changed when the p-FFC formed, the distance data was fit to a model of one near and one far Gaussian functions (Fig. S11; Table S5). Shifts in population between the near and far components were most prominent for the 370-658, 490-658 and 490- 746 variants, which contain sites involved in core clock interactions (Fig. S11). The proportion of the closer component increased from ∼30% to ∼70% in the 370-658 variant (Fig. S11) and from ∼10% to ∼70% in the 490-746 variant (Fig. 5F) when bound to FRH and CK1. Similar trends were observed when we compared the percentage change of the average separation observed in FRQ alone versus when it is bound to FRH and CK1 (Fig. 5H). The 490, 658 and 746 sites reside in regions important for WCC interactions (Guo, et al., 2010) and FRH may organize such binding determinants. Notably, the 370SL:746SL spin pair showed little dipolar coupling in free p-FRQ but resolved into an observable mid-range separation in the presence of CK1 and FRH (Fig. 5C), perhaps because position 746 is within the FRH binding region. Thus, CK1 and FRH order p-FRQ to an appreciable degree and the two enzymes are adjacently positioned within the conformationally heterogeneous complex.

### FRQ exhibits behavior consistent with LLPS *in vitro* and *in vivo*

The unstructured regions and multivalent interactions of IDPs can lead to liquid-liquid phase separation (LLPS) (Martin, E. W., et al., 2020; Martin, E. W., et al., 2020). Such de-mixing often manifests in the formation of biomolecular condensates and membrane-less organelles (MLOs) within cells (Posey, et al., 2018). As has been previously noted (Pelham, et al., 2020), FRQ may be capable of LLPS owing to its extensive disorder. We found that FRQ has a high propensity to undergo LLPS based on a set of computational tools that evaluate charge distribution and clustering, long-range aromatic and non-aromatic π:π contacts and nucleic acid binding propensities (Bolognesi, et al., 2016; Vernon, et al., 2018). The overall FRQ PScore, which grades LLPS propensity on these properties (Vernon, et al., 2018), was 6.39, with three regions scoring above a value of 2, the threshold for favorable LLPS (Fig. S12). The region with the highest PScore also accounted for the majority of LLPS propensity as determined by the catGRANULE prediction algorithm (Bolognesi, et al., 2016). Furthermore, we found that these methods also predicted the functional analogs dPER and hPER1 to phase separate (Fig. 6A, Fig. S13). The FRQ functional analogs share physicochemical characteristics with FRQ, particularly an enrichment of hydrophilic residues, increases in Gly and Pro content and a depletion of hydrophobic residues, even though their positional sequence conservation with FRQ is low (Fig. S14 and S15; see also (Jankowski, et al., 2022; Pelham, et al., 2021)). Similar amino acid composition patterns among FRQ, dPER and hPER1, include the most common residues in each protein and its UniRef50 cluster (sequences with at least 50% identity with the target protein) being serine, proline and glycine (∼30% of total sequence) accompanied by a high depletion in aromatic residues (only ∼7.5%) (Fig. S15). In addition, all of the sequences have a reduced proportion of hydrophobic residues such as A, I, L, and V and a slightly acidic isoelectric point. Thus, FRQ, dPER and hPER1 all have low overall hydropathy (Fig. S14). However, in addition, FRQ and its homologs are unusual in their high Arg and Asp content, which tend to separate in charge blocks throughout their sequences (as indicated by high kappa values, Fig. S15; see also (Jankowski, et al., 2022)). Such charge clustering correlates with LLPS (Somjee, et al., 2020). Overall, this analysis suggests that like intrinsic disorder, LLPS driven by the physiochemical properties of the constituent residues, may be a conserved feature in the negative arm of the circadian clock.

**Figure 6:**
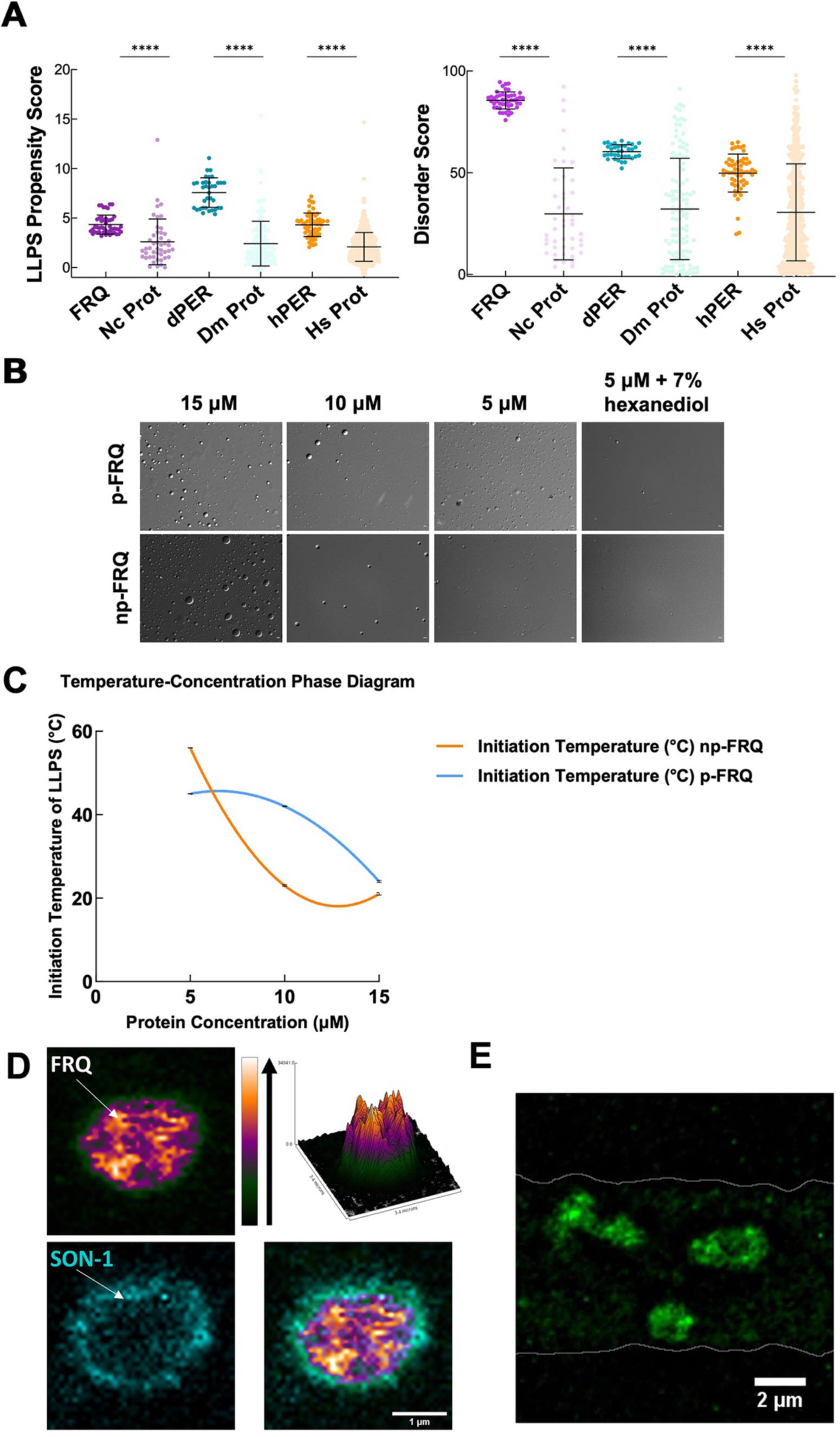
FRQ undergoes LLPS in agreement with predictions from its sequence properties and those of its functional analogs. (A) LLPS propensity predictions using “Pi-Pi” and IUPred2A disorder predictions are shown for FRQ (purple), dPER1 (teal) and hPER1 (orange) as compared to values for proteins of similar length from their respective proteomes (Prot): *Neurospora crassa*, Nc; *Drosophila melanogaster*, Dc; *Homo sapiens*, Hc. The mean and standard deviations are represented as horizontal and vertical black bars, respectively. Note: **** denotes a p-value <0.0001 obtained from a Mann-Whitney U-test (B) DIC microscopy images of various concentrations of p-FRQ (in 150 mM NaCl, 25 mM Tris pH 8.0) and np-FRQ (in 500 mM NaCl, 25 mM Tris pH 8.0) at 25°C; images shown at 100x magnification. Scale bar = 2μm (C) Temperature vs concentration phase diagram derived from the results of the turbidity assays shown in Fig. S17. Both p-FRQ and np-FRQ undergo an LCST phase transition on or above the line (a second order polynomial fit to the data points). The error bars of each point (about the size of the points) reflect the 95% confidence interval of the mean. The phase boundary for p-FRQ represents the transition apparent in the temperature scans of Fig. S17. Some p-FRQ phase separation already occurs at low temperature under these buffer conditions and thus the region below the line does not represent fully soluble p-FRQ. (D) Live cell fluorescent imaging of FRQ^mNeonGreen^ in nuclei of *Neurospora* hyphae. Upper left panel shows the FRQ channel, represented as a multicolor heat map of fluorescence intensity; upper right panel shows a surface plot derived from the raw FRQ image to emphasize regions of concentration; lower left shows nucleoporin SON-1^mApple^, which localizes to the cytoplasmic face of the nuclear envelope; bottom right shows the FRQ:SON-1 merge image. The images were acquired on a Zeiss 880 laser scanning confocal microscope and were smoothed by bicubic interpolation during 10-fold enlargement from 42 X 42 pixels to 420 X 420 pixels. (E) FRQ[mNeonGreen] nuclear foci shown within an *N. crassa* syncytium mycelium (outlined by white lines), cropped from movie S1. Scale bar = 2μm.

We tested whether both np-FRQ and p-FRQ can undergo LLPS *in vitro.* In each case, upon exchanging the purified protein into their respective phase separation buffers (which lack any crowding agents), we observed the appearance of a turbid solution that contained microscopic droplets (Fig. 6B). These droplets scaled in size and number with increasing protein concentration, dissolved upon the addition of 1,6-hexanediol, and were shown to dock and fuse to one another, thereby confirming their liquid-like properties (Fig. 6B, Fig. S17). UV-Vis turbidity assays as a function of temperature revealed that FRQ undergoes a low critical solution temperature (LCST) phase transition that is concentration and phosphorylation status-dependent. (Fig. 6C; Fig. S17,Tables S6-7). As expected, phase separation increased as temperature and protein concentration increased as indicated by growing amplitudes and earlier phase separation initiation temperatures (Fig. 6C; Fig. S17; Table S6-7). Lower salt concentrations favored the LLPS of p-FRQ, whereas higher salt disfavored LLPS, compared to higher salt concentrations favoring LLPS for np-FRQ (Fig. S17). Although the NaCl concentration for np-FRQ LLPS conditions is non-physiological *per se*, the ionic strength conditions and effective solute concentrations in cells are comparatively high (Liu, B. Q., et al., 2017). Furthermore, the higher absorbance values of p-FRQ at lower temperatures relative to that of np-FRQ indicates complex phase behavior for the phosphorylated isoform that cannot adequately be explained by an LCST phase transition alone (Fig. S17). At temperatures below the approximated phase boundary indicated by the temperature transitions (Fig. 6C; Fig. S17) some p-FRQ phase separation has already occurred in the phase-separation buffer. Thus, FRQ phosphorylation alters its behavior consistent with LLPS as evidenced by its non-zero, concentration-dependent absorbance at lower temperatures, mostly likely by increasing the relative negative charges, changing the charge patterns, and providing more sites for multivalent interactions within the molecule (Fig. 6C, Fig. S16-17)(Somjee, et al., 2020; Szabó, et al., 2022).

Live cell imaging of *Neurospora crassa* expressing FRQ tagged with the fluorescent protein mNeonGreen at its C-terminus (FRQ^mNeonGreen^) revealed heterogeneous patterning of FRQ in nuclei (Fig. 6D,E). FRQ thus tagged and driven by its own promoter is expressed at physiologically normal levels, and strains bearing FRQ^mNeonGreen^ as the only source of FRQ are robustly rhythmic with a slightly longer than normal period length. Live-cell imaging in *Neurospora crassa* offers atypical challenges because the mycelia grow as syncytia, with continuous rapid nuclei motion during the time of imaging. This constant movement of nuclei is compounded by the very low intranuclear abundance of FRQ and the small size of fungal nuclei, making not readily feasible visualization of intranuclear droplet fission/fusion cycles or intranuclear fluorescent photobleaching recovery experiments (FRAP) that could report on liquid-like properties. Nonetheless, bright and dynamic foci-like spots were observed well inside the nucleus and near the nuclear periphery, which is delineated by the cytoplasm-facing nucleoporin Son-1 tagged with mApple at its C-terminus (Fig. 6D,E, Movie S1). Such foci are characteristic of phase separated IDPs (Bartholomai, et al., 2022; Caragliano, et al., 2022; Gonzalez, A., et al., 2021; Tatavosian, et al., 2019) and share similar patterning to that seen for clock proteins in *Drosophila* (Meyer, et al., 2006; Xiao, et al., 2021), although the foci we observed are substantially more dynamic than those reported in Drosophila.

### Phase separated FRQ recruits FRH and CK1 and alters CK1 activity

The PScore and catGRANULE LLPS predictors gave low overall scores for the FRQ partners, FRH and CK1 (Fig. S13). This result is unsurprising because these proteins consis of predominantly folded domains. Nonetheless, FRH and CK1 may still act as clients that localize to the FRQ-formed LLPS-like droplets. We chose to investigate whether FRH and CK1 could be recruited into droplets with fluorescent microscopy. Structured proteins are not well tolerated in the temperature-dependent turbidity assays because of the large signal that they give upon domain unfolding. We found that fluorescently-labeled FRH and CK1 were recruited into phase separated FRQ droplets, but a control protein labeled with the same fluorophore, *E. coli* CheY, was not (Fig. 7A). The ability of FRQ to recruit its partner proteins into phase separated droplets provides a means to sequester them and potentially control their availabilities and activities.

**Figure 7:**
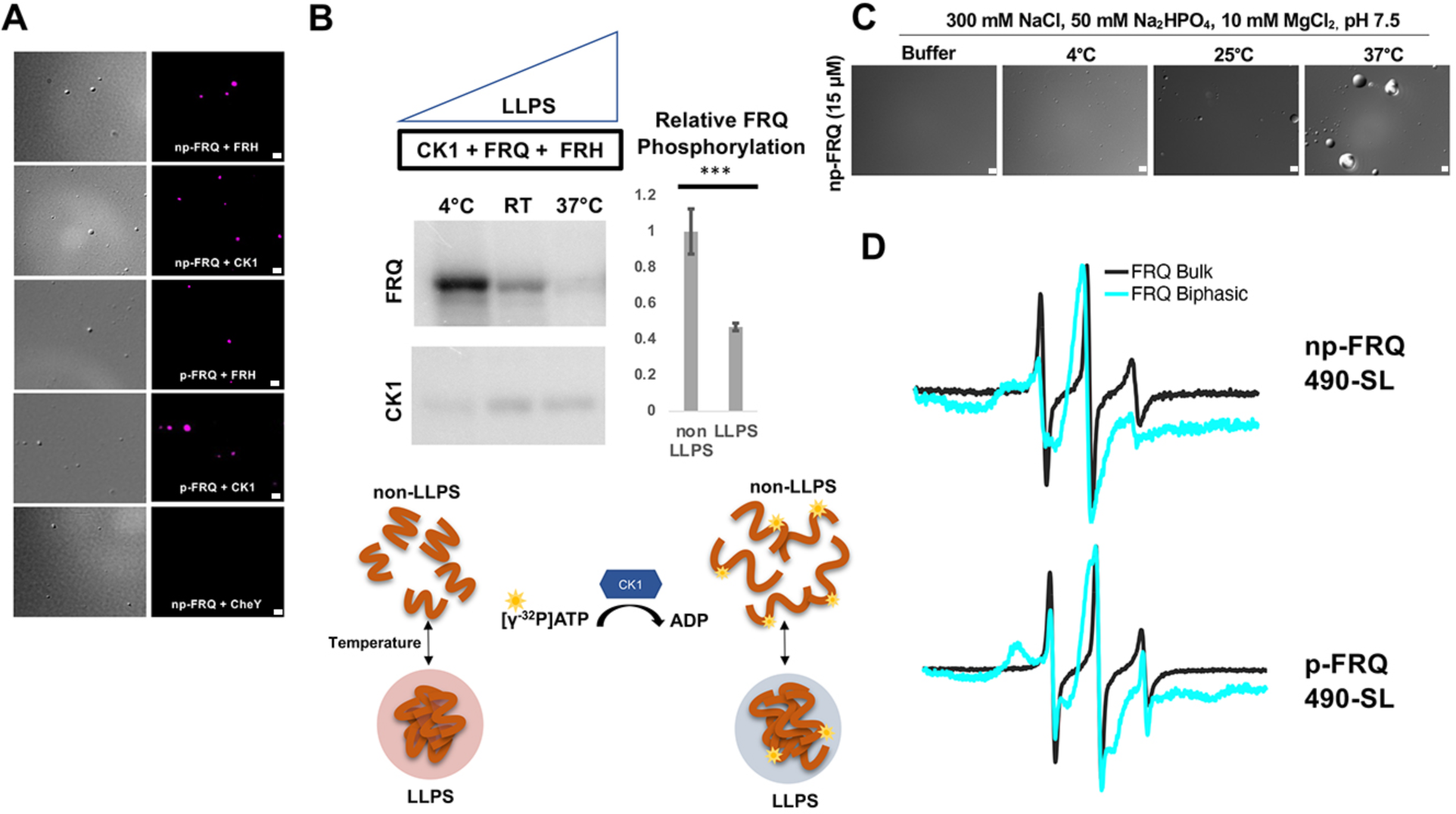
Conditions favoring LLPS alter the structural and enzymatic properties of the FFC. (A) Differential Interference Contrast (DIC) and fluorescence microscopy images at 25 °C of phase separated np-FRQ or p-FRQ (5 μM each) droplets in 500 mM KCl, 20mM Na_2_HPO_4_, pH 7.4 with either equimolar Cy5-labeled FRH, CK1 or CheY (control) visualized using Cy5 fluorescence. Scale bar = 2μm . (B) (Top) Autoradiography of FRQ (20 μM) (left) phosphorylated by CK1 in the presence of FRH at increasing temperatures. (Bottom) Schematic showing the nature and results of the autoradiography assay depicted above. The FRQ phosphorylation under LLPS conditions was reduced relative to non-LLPS conditions. The complete autoradiograms are shown in Fig. S13. Quantification of FRQ phosphorylation by CK1 at RT under LLPS and non-LLPS conditions with phosphorylation levels under non-LLPS conditions normalized to 1. Note: *** denotes a p-value << 0.05 obtained from a student’s t-test. (C) DIC microscopy images of 15 μM phase separated np-FRQ under the same conditions (i.e. buffer and temperature) as the phosphorylation assay shown in (B). Scale bar = 2μm . (D) X- band (9.8GHz) RT cw-ESR spectra of FRQ labeled with MTSL at the 490 site in solubilizing buffer (black) and under conditions that promote LLPS (cyan). Data represented as first-derivative spectra with the horizontal axes depicting the static field strength (typically 3330 to 3400 G), and the vertical axis depicting the change in magnetic susceptibility with respect to the static field, in arbitrary units.

LLPS has been suggested to modulate the activities of enzymes by creating a microenvironment to alter substrate sequestration and product release (Shapiro, et al., 2021). We tested the effect of FRQ LLPS on its phosphorylation by CK1, a key reaction for regulating the period of the clock (Liu, X., et al., 2019). We compared the phosphorylation of FRQ by CK1 in a buffer that supports phase separation under different temperatures, using the latter as a means to control the degree of LLPS without altering the solution composition. Both the UV-Vis turbidity assays as well as the DIC micrographs of the protein at different temperatures establish that FRQ LLPS increases substantially with temperature (Fig. 6C; Fig.7C). Furthermore, LLPS of p-FRQ and np-FRQ both depend highly on buffer composition (Fig. S17). CK1 autophosphorylation served as an internal control and its temperature dependence was consistent with previous studies that show increased catalytic activity at higher temperatures (Marzoll, et al., 2022). As temperature increases drove more LLPS, the phosphorylation of FRQ decreased, even as CK1 autophosphorylation increased (Fig. 7B; Fig. 8). Quantitatively, we observed a reduction in FRQ phosphorylation by two-fold under LLPS conditions relative to conditions under which it does not phase separate (Fig. 7B).

**Figure 8:**
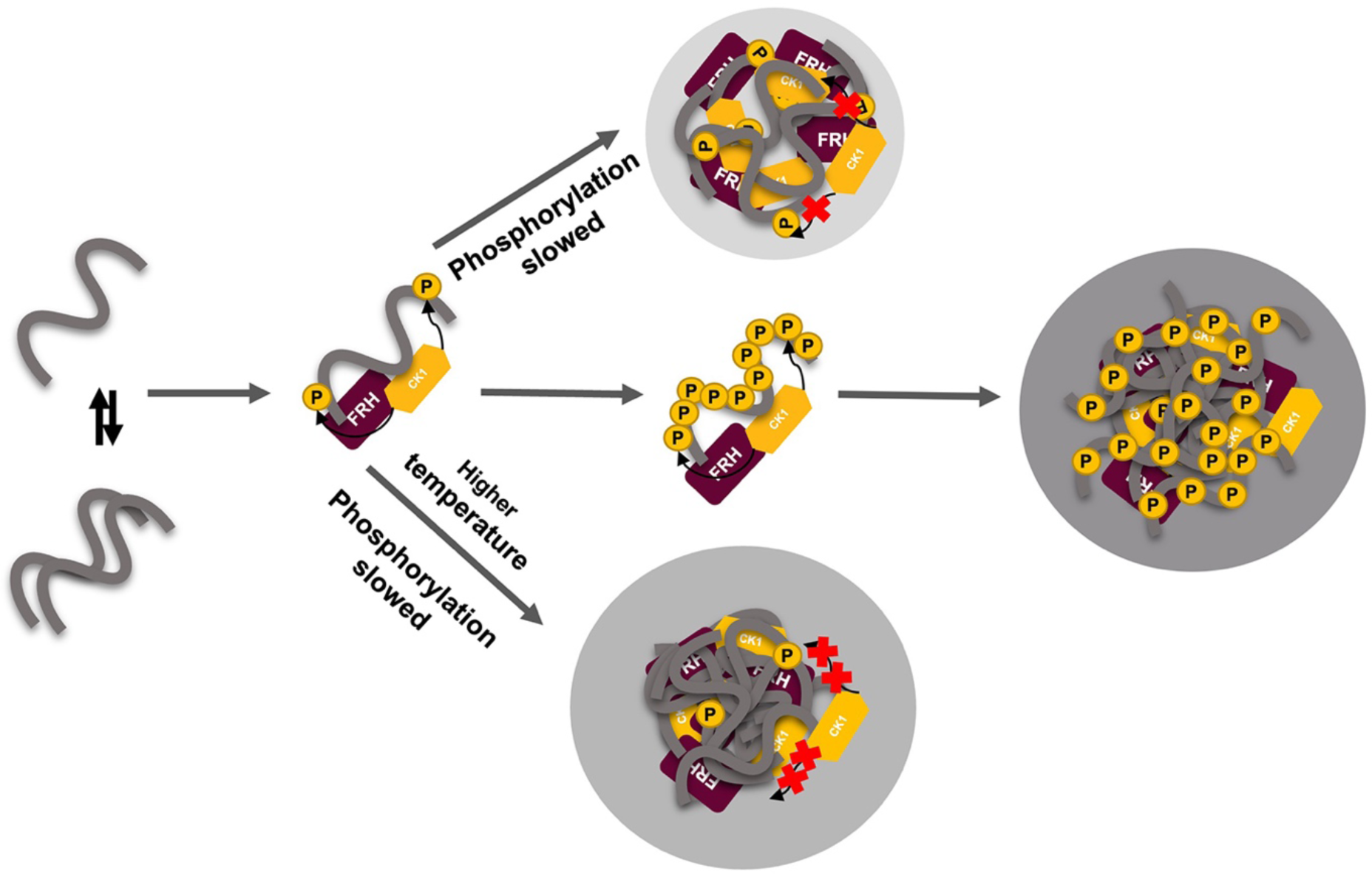
LLPS may play a role in temperature compensation of the clock through modulation of FRQ phosphorylation. Reduced CK1 phosphorylation of FRQ causes both longer periods (Mehra, et al., 2009) and loss of temperature compensation (manifested as a shortening of period at higher temperature) (Hu, et al., 2021; Liu, X., et al., 2019). Thus, the ability of increased LLPS at elevated temperature (larger grey circle) to reduce FRQ phosphorylation by CK1 will counter a shortening period that would otherwise manifest in an under compensated system. As further negative feedback, LLPS is also promoted by increased FRQ phosphorylation, which in turn will reduce phosphorylation by CK1. Thus, both increased FRQ phosphorylation and temperature favor LLPS and reduction of CK1 activity.

Through application of cw-ESR, we probed FRQ dynamics when it undergoes LLPS . We observed a striking rigidification of spin-labeled FRQ when LLPS was promoted by buffer conditions (Fig. 7D). The reduction of CK1-mediated FRQ phosphorylation, which likely requires substantial dynamics given the distribution of phosphorylation sites throughout the FRQ sequence, may reflect this change in molecular dynamics. Thus, in addition to affecting the localization of FRQ and its partners, LLPS also impacts the enzymatic activity of the complex, by perhaps modulating molecular encounters and motions essential for large scale phosphorylation.

## Discussion

Within the TTFLs that define eukaryotic circadian clocks, large repressor proteins like FRQ compose cellular timing mechanisms by coordinating transient protein-protein interactions, nuclear entry, targeted protein degradation, and transcriptional repression (Pelham, et al., 2020). They compose dynamic macromolecular assemblies that achieve specificity in their functions despite assuming highly variable structures (Pelham, et al., 2020). Our purification scheme enabled a multi-pronged biophysical characterization of FRQ. The majority of previous such studies on IDPs, particularly those that undergo de-mixing, have involved truncated proteins, excised domains or peptides of sizes less than 200 residues (Babinchak, et al., 2019; Emmanouilidis, et al., 2021; Lin, et al., 2021; Lin, et al., 2019; Seal, et al., 2021). FRQ represents a full-length, nearly 1000-residue IDP that can be investigated with respect to post-translational modification, interaction partners and phase separation. As anticipated from its sequence (Hurley, et al., 2013; Pelham, et al., 2020), this direct characterization of structural properties shows that FRQ is indeed a highly dynamic protein, but even in isolation, it is not completely disordered. The SAXS and ESR data indicate that FRQ has compact features that localize to an ordered core interspersed with dynamic regions. Notably the shape of the Kratky plots generated from the SAXS data suggest a degree of disorder that is substantially greater than that expected of a molten globule (Kataoka, et al., 1997), but far from that of a completely denatured protein (Kikhney, et al., 2015; Martin, Erik W., et al., 2021). Similarly, the DEER distributions, though non-uniform across the various sites examined, indicate more disorder than that of a molten globule (Selmke, et al., 2018) but more order than a completely unfolded protein (van Son, et al., 2015). Although FRQ lacks the well-defined PAS domains of the analogous PER proteins (Pelham, et al., 2020), it also maintains a structured core to anchor its more dynamic elements. The regions that bind CK1 and FRQ are the most ordered, and they organize the two partners beside each other; in fact, they are in remarkably close proximity, as judged by spin-labels in their ATP binding pockets or in their binding domains. FRH mediates contacts of the FFC to the WCC (Conrad, et al., 2016), whereas CK1 phosphorylates the WCC to close the TTFL (Cha, et al., 2014; Larrondo, et al., 2015; Liu, X., et al., 2019; Wang, et al., 2019); hence, the targeting of CK1 to the WCC likely depends on FRQ associating FRH with CK1 (Wang, et al., 2019).

Previous proteolytic sensitivity studies have suggested that increased phosphorylation across the day transitions FRQ from a “closed” to a more “open” state (Liu, X., et al., 2019; Pelham, et al., 2020; Querfurth, et al., 2011). Our data paint a more nuanced picture. The SAXS data reveal that p-FRQ and p-FFC are more expansive and show greater flexibility than do np-FRQ and np-FFC, respectively. Furthermore, np-FRQ has a greater tendency to self-associate. In the cell, completely unphosphorylated FRQ is likely short-lived, if present at all (Baker, et al., 2009). Phosphorylation may stabilize FRQ against aggregation, yet also provides for enough conformational heterogeneity to poise FRQ near a degradation threshold (Larrondo, et al., 2015). This idea is supported by the cw-ESR data (Fig. 3), which shows that phosphorylation causes non-uniform local structural changes with some regions increasing, while others decreasing, in flexibility. The DEER data also demonstrates that binding to FRH and CK1 increases compactness of p-FRQ, particularly in the central-core that is responsible for binding the partners. In contrast, the FRQ C-terminus gains considerable flexibility in the FFC. FRQ has been suggested to assume a dimeric state, because of a predicted coiled-coil region and formation of mixed dimers formed in cellular pull-down assays with differentially tagged FRQ subunits (Cheng, et al., 2001). Np-FRQ does indeed dimerize when isolated, whereas p-FRQ is primarily monomeric alone. Furthermore, SAXS data indicates there is one copy of FRQ in both the reconstituted np-FFC and p-FFC. Hence phosphorylation has the potential to regulate the FRQ oligomeric state, which may in turn be linked to FFC assembly. Despite being a monomer in isolation, the pull-down data (Cheng, et al., 2001) indicate that association of p-FRQ may be mediated by other cellular factors, including LLPS.

FRQ has physicochemical features embedded within its sequence that give rise to LLPS and the FRQ clock analogs exhibit similar characteristics (Fig. S13-15). FRQ and negative-arm proteins in general belong to the “flexible disorder” class of IDPs (Shapiro, et al., 2021), wherein amino acid patterns that generate intrinsic disorder are conserved, but amino acid sequence per se, is not (Bellay, et al., 2011). Moreover, although these proteins all clearly contain regions of low-complexity sequence (Hurley, et al., 2013; Marzoll, et al., 2022; Pelham, et al., 2020), their conserved features include not only their intrinsic disorder, but also their propensity for LLPS (Fig. 6A,Fig. S13). These two properties, although related, are not directly correlated (Bolognesi, et al., 2016; Martin, E. W., et al., 2020; Szabó, et al., 2022; Vernon, et al., 2018). In particular, a preponderance of aromatic and non-aromatic π-π planar contacts and charge-dense or charge- repeat regions drive LLPS (Szabó, et al., 2022; Vernon, et al., 2018) and these features are consistently found in clock repressors, particularly FRQ. The similar amino acid compositions of these proteins may then potentially give rise to their shared function. Moreover, extensive phosphorylation may alter LLPS propensity, because it increases extended states, solvation of the peptide backbone, charge density and intermolecular hydrogen bonding capacity (Bolognesi, et al., 2016; Szabó, et al., 2022; Vernon, et al., 2018). For FRQ, the sequence predictions are borne out experimentally, with p-FRQ displaying a different phase behavior relative to np-FRQ. In addition to being more soluble, dynamic and less aggregated than np-FRQ, p-FRQ also phase separates at lower temperatures and under lower salt conditions relative to np-FRQ. Increased salt disfavors p-FRQ phase separation, which when compared to np-FRQ, is perhaps consistent with more electrostatically driven de-mixing due to the high degree of phosphorylation.

Live cell imaging of fluorescently-tagged FRQ proteins is consistent with FRQ phase separation in *N. crassa* nuclei. FRQ is plainly not homogenously dispersed within nuclei, and the concentrated foci observed at specific positions in the nuclei indicate condensate behavior similar to that observed for other phase separating proteins (Bartholomai, et al., 2022; Caragliano, et al., 2022; Gonzalez, A., et al., 2021; Tatavosian, et al., 2019; Xiao, et al., 2021). While ongoing experiments are exploring more deeply the spatiotemporal dynamics of FRQ condensates in nuclei, the small size of fungal nuclei as well as their rapid movement with cytoplasmic bulk flow through the hyphal syncytium makes these experiments difficult. Of particular interest is drawing comparisons between FRQ and the Drosophila Period protein, which has been observed in similar foci that change in size and subnuclear localization throughout the circadian cycle (Meyer, et al., 2006; Xiao, et al., 2021), although it must be noted that the foci we observed are considerably more dynamic in size and shape than those reported for PER in Drosophila (Xiao, et al., 2021). A very recent manuscript (Xie, et al., 2024) calls into question the importance and very existence of LLPS of clock proteins at least in regards to mammalian cells, noting that it may be an artifact of overexpression in some instances where it is seen, and that at normal levels of expression there is no evidence for elevated levels at the nuclear periphery. Artifacts resulting from overexpression are unlikely to be a problem for our study and that of Xiao et al as in both cases clock proteins were tagged at their endogenous locus and expressed from their native promoters. Although we noted enrichment of FRQ^mNeonGreen^ near the nuclear envelope in our live-cell imaging, there remained abundant FRQ within the core of the nucleus.

Phosphorylation is known to modulate LLPS in other systems, but the effects are variable. For example, phosphorylation of the transcriptional silencing factor HP1α also enhances LLPS, whereas only the non-phosphorylated form of Xenopus CPEB4 involved in translation of mRNA poly-A tails undergoes LLPS (Larson, et al., 2017; Seal, et al., 2021). FRQ, especially p-FRQ, may achieve its inhibitory function by sequestering the FFC and WCC into membraneless organelles. The large, disordered domains and capacity for multivalency of the WC proteins (Pelham, et al., 2020) may also facilitate their partitioning into such condensates. In the fly clock, PER and the WCC analog CLK produce liquid-like foci in the nuclei that are most prevalent during the repressive phase of the circadian cycle (Xiao, et al., 2021). These large foci, which are few and localized to the nuclear periphery, may serve to organize clock-controlled genes during repression (Xiao, et al., 2021). LLPS may also influence the nuclear transport of FRQ by affecting interactions with the nuclear import machinery as has been observed with proteins such as TD43 and FUS (Gasset-Rosa, et al., 2019; Gonzalez, Abner, et al., 2021). Conditions favoring LLPS reduce the ability of FRQ to act as a substrate for CK1 (Fig. 7B), and this reactivity correlates with reduced dynamics of FRQ in the phase separated state (Fig. 7D). Domains and disordered regions from other spin-labeled proteins show similar reduced dynamics when induced to phase separate and monitored by cw-ESR spectroscopy (Babinchak, et al., 2019; Emmanouilidis, et al., 2021; Lin, et al., 2021; Seal, et al., 2021), although the temperature and salt dependencies of the ordering behavior can vary considerably, with some species showing little change upon LLPS (Lin, et al., 2019).

In the cell, the impact of LLPS on CK1 activity may modulate the timing of FRQ phosphorylation, which is closely linked to clock period (Larrondo, et al., 2015; Marzoll, et al., 2022). Moreover, and collectively, the data from Fig. 6 and 7 show that as temperature increases, LLPS-like behavior increases, and as LLPS increases, the ability of CK1 to phosphorylate FRQ decreases. We believe that the reduced CK1 kinase activity toward FRQ as a substrate is directly due to the impact of the generated LLPS milieu, i.e. the changes in structural/dynamic properties of FRQ and/or CK1 induced by the effects of being a phase separate microenvironment, which could be substantially different from non-phase separated buffer environment. For example, previous work done on the disordered region of DDX4 (Brady, et al., 2017; Nott, et al., 2015) show that even the amount of water content and stability of biomolecules such as double strand nucleic acids encapsulated within the droplets differ between non- and phase separated DDX4 samples. Indeed, the spin-labeling experiments indicate that the dynamics of FRQ have been altered by LLPS (Fig. 7D).

The rate of FRQ phosphorylation dictates period length (Larrondo, et al., 2015). Therefore, this negative feedback of temperature on period mediated by LLPS formation could provide a mechanism for circadian temperature compensation of period length (Fig. 8). Sequestration of CK1 (and FRH) into condensates may also render these factors unavailable to target other cellular substrates. Phosphorylation alters FRQ interactions throughout the circadian cycle, not unlike the workings of the cyanobacterial clock, which relies upon a timed phospho-code to regulate enzymatic components (Rust, et al., 2007; Wang, et al., 2019). We demonstrate that FRQ likewise responds to phosphorylation with non-uniform conformational changes across its sequence and an increased propensity for LLPS. It is the latter property that appears conserved among negative-arm repressor proteins in eukaryotic clocks and may thus be a critical function of their intrinsic disorder and extensive post-translational modifications.

## Materials and Methods

FRQ was obtained by recombinant expression in *E. coli* and purification under denaturing conditions followed by removal of denaturant. FRH and CK1 were produced using more standard protocols. SEC-MALS was performed with Wyatt Technologies DAWN HELEOS-II multi-angle light scattering detector and SEC-SAXS was carried out at the Cornell Synchrotron G1 station. Protein labeling for ESR and microscopy experiments made use of enzymatic labeling and reactive cysteine residue probes. Cw-ESR spectroscopy was carried out on a Bruker E500 ESR spectrometer operating at X-band, whereas DEER spectroscopy utilized a four-pulse protocol on a Bruker E580 spectrometer operating at Q band. Turbidity was as measured by scattering at 300 - 600 nm under different buffer conditions and a Peltier-driven temperature ramp. Microscopy was carried out on a Zeiss Axio Imager Upright Trinocular Fluorescence Microscope at 100x magnification. Phosphorylation assays involved reaction with [γ-^32^P] ATP, protein band resolution by SDS-PAGE and phosphorimage analysis. Mass-spectrometry and phospho-peptide detection was carried out on tryptic peptides analyzed by MS/MS with a CID ion-trap. *Neurospora crassa* strain generation targeted the endogenous *frq* locus with DNA encoding *N. crassa* codon-optimized mNeonGreen appended to the C-terminus of the *frq* ORF. For live-cell imaging a section near the edge of a growing mycelium was removed and inverted on a chambered cover glass. Images were collected with 488 nm laser excitation using a laser scanning confocal microscope equipped with a spinning disk scan head and a CMOS camera. Detailed descriptions of the protocols can be found in the *Supplementary Materials and Methods*.

Author contributions

D.T., N,M., B.M.B., J.C.D., A.B. and B.R.C. designed research; D.T., N.M., B.M.B., and S.C. performed research; D.T., N,M., B.M.B., S.C., J.C.D., A.B. and B.R.C. analyzed data; D.T. and B.R.C. wrote the paper with contributions and editing from all authors.

The authors declare no competing interests.

## Data Availability

The mass spectrometry FRQ phosphorylation data have been deposited to the ProteomeXchange Consortium via the PRIDE partner repository with the dataset identifier PXD037938. All other data are included in the article and/or supporting Information.

## Supporting information

which is delineated by the cytoplasm-facing nucleoporin Son-1 tagged with mApple at its C-terminus (Fig. 6D,E, Movie S1).

## Acknowledgements.

We thank the Cornell High Energy Synchrotron Source (CHESS) and the National Biomedical Center for Advanced ESR Technologies (ACERT) for access to data collection facilities. This work was supported by grants from the National Institutes of Health: R35GM122535 to BRC, R35GM118021 to JCD; R35GM138097 to AB, 1F31GM143890 to NM, and T32-008704 to BMB and from The Pew Charitable Trusts to AB. ACERT is supported by P41GM103521 and R24GM146107. CHESS is supported by NSF award DMR-1332208 and NIH/NIGMS award P30GM103485.

## Supplementary Information

### Materials and Methods

#### Protein Expression and Purification

Full length L-FRQ (Uniprot ID:P19970), CK1 (CK1a; Uniprot ID:V5IR38) and FRHΔN (residues 100-1106) (Uniprot ID:Q1K502) were sub-cloned into a pET28a vector (Novagen) bearing an N-terminal His_6_-tag and kanamycin resistance. Note: L-FRQ contained the S513A substitution to improve protein stability as noted in (Querfurth, C., et al., 2007). The plasmids encoding the proteins were transformed into BL21 (DE3) *E. coli* cells, grown in LB media containing 50 μg ml^−1^ kanamycin to an optical density (OD_600_) of ∼0.6–0.8 at 37 °C and protein expression was induced with 1 mM IPTG at 17 °C or room temperature. Cells were harvested after 12–18 h by centrifugation at 5,000 rpm and 4 °C. FRH and CK1 were purified by standard Ni-NTA chromatography followed by ion-exchange and size-exclusion chromatography. Briefly, cells were resuspended in lysis buffer (50 mM Tris pH 8.0, 500 mM NaCl, 10% glycerol, 1 mM MgCl_2_, 5 mM imidazole) and sonicated on ice. The lysate was clarified by centrifugation at 20,000 rpm for 1 hr. The clarified lysate was applied to a gravity Ni-NTA column and washed with 5 column volumes (CVs) of wash buffer (50 mM Tris pH 8.0, 500 mM NaCl, 10% glycerol, 1 mM MgCl_2_, 20 mM imidazole.) The protein was eluted using the elution buffer (50 mM Tris pH 8.0, 500 mM NaCl, 10% glycerol, 1 mM MgCl_2_, 500 mM imidazole). The protein was then diluted into IEX (ion-exchange) bufferA (50 mM HEPES pH 7.5, 10% glycerol), applied to an S cation exchange column (GE Healthcare) and eluted over a 20 CV gradient against IEX bufferB (50 mM HEPES pH 7.5, 1 M NaCl, 10% glycerol). Fractions containing the protein were pooled, concentrated, and run over an S200 26/60 gel-filtration column pre-equilibrated in gel-filtration buffer (GFB; 50 mM HEPES, 200 mM NaCl, 10% glycerol.) L-FRQ was purified under denaturing conditions. Cells were resuspended in lysis buffer (50 mM Na_2_PO_4_ pH 7.42, 300 mM NaCl, 5 mM imidazole, 4 M guanidinium hydrochloride (GuHCl)) and sonicated on ice. The lysate was clarified by centrifugation at 20,000 rpm for 1 hr. The clarified lysate was applied to a gravity Ni-NTA column and washed with 3 CV of lysis buffer. The protein was eluted using the elution buffer (50 mM Na_2_PO_4_ pH 7.42, 300 mM NaCl, 500 mM imidazole, 4 M GuHCl). The protein was concentrated and applied to a S200 26/60 gel-filtration column pre-equilibrated in 50 mM Na_2_PO_4_ pH 7.42, 300 mM NaCl, 5mM Imidazole, 4 M GuHCl. Fractions containing the protein were pooled and concentrated. The concentrated protein was flash-frozen and stored at -80 °C. Prior to experiments, the purified L-FRQ was exchanged into a non-denaturing buffer (50 mM Na_2_PO_4_ pH 7.42, 300 mM NaCl, 200 mM arginine, 10% glycerol). For FFC preparations, GFB (50 mM HEPES pH 7.42, 200 mM NaCl, 10% glycerol) was used.

#### SEC-MALS

Protein samples between 50-75 μM were injected onto a Superose-6 (GE Lifesciences), equilibrated with 50 mM Na_2_PO_4_ pH 7.42, 300 mM NaCl, 200 mM arginine, 10% glycerol. The gel filtration column was coupled to a static 18-angle light scattering detector (DAWN HELEOS-II) and a refractive index detector (Optilab T-rEX) (Wyatt Technology, Goleta, CA). Data were collected every second at a flow rate of 0.5 mL/min and analysis was carried out using ASTRA VI, yielding the molar mass and mass distribution (polydispersity) of the sample. The monomeric fraction of BSA (Sigma, St. Louis, MO) was used to standardize the light scattering detectors and optimize data quality control.

#### SEC-SAXS

X-ray scattering experiments were performed at the Cornell High Energy Synchrotron Source (CHESS) G1 station. Size-exclusion chromatography-coupled SAXS (SEC–SAXS) experiments were performed using a Superdex 200 5/150 GL or Superose-6 5/150 GL (3 mL) column operated by a GE AKTA Purifier at 4 °C with the elution flowing directly into an in-vacuum X-ray sample cell. Samples were centrifuged at 14,000×*g* for 10 min at 4 °C before loading onto a column pre-equilibrated in a matched buffer (50 mM Na_2_PO_4_ pH 7.42, 300 mM NaCl, 200 mM arginine, 10% glycerol). Samples were eluted at flow rates of 0.05–0.1 mL min^−1^. For each sample, 2-s exposures were collected throughout elution until the elution profile had returned to buffer baseline, and scattering profiles of the elution buffer were averaged to produce a background-subtracted SEC–SAXS dataset. The scattering data was processed in RAW (Hopkins, et al., 2017) and regions with homogenous R_G_ values were chosen for the Guinier and Kratky analyses.

#### Protein Labeling for ESR Spectroscopy and Microscopy Experiments

Sort/AEP1-tagged FRQ was cloned into a pET28a (Novagen) vector as described in (Chandrasekaran, et al., 2021). The production of the SortaseA and AEP1 enzymes and their associated peptides (GGGGC for SortaseA and CNGL for AEP1) is also described in (Chandrasekaran, et al., 2021) and (Nguyen, et al., 2015), respectively. To enzymatically label FRQ with either SortaseA or AEP1, 1:1 protein and SortaseA/AEP1 as well as 2x excess peptide were incubated overnight to allow the reaction to reach completion. The following day, the reaction was run on a Superpose 6 10/300 analytical SEC column to remove unreacted peptide and enzyme, and the protein containing fractions were pooled and concentrated. To prepare the ADP- β-S-SL samples, ADP-β-S-SL was synthesized as previously reported in (Muok, et al., 2018). 100 μL of buffer containing 100 μM of ADP-β-S-SL and 100 μM FRH or CK1 were incubated overnight and then washed with GFB (50 mM HEPES pH 7.42, 200 mM NaCl, 10% glycerol) through 5x dilution followed by re-concentration to remove any unbound ADP-β-S-SL. 30 μL of each sample were used for data collection. For MTSL labeling, the protein was incubated with 5x molar excess of MTSL overnight to allow the reaction to reach completion. The following day, the reaction was run on HiTrap Desalting column (GE Life Sciences) to remove any excess spin-label.

To prepare fluorophore labeled FRH and CK1, the purified proteins were incubated with 2x molar excess Cy5-maleimide dye (GE Life Sciences) overnight to label any accessible cysteines on the target proteins. The excess dye was removed by running the reaction over a HiTrap desalting column equilibrated with GFB (50 mM HEPES pH 7.42, 200 mM NaCl, 10% glycerol).

#### ESR Spectroscopy

The cw-ESR spectra were obtained on a Bruker E500 ESR spectrometer operating at X-band (∼9.4 GHz). The ESR measurements were carried out at room temperature with a modulation amplitude of 0.5-2 G and modulation frequency of 100 kHz. All spectra were digitized to 1024 points with a sweep width of ∼100 G and an average of 4–16 scans. The δ values were calculated using the display distance function on the Bruker Xepr 2.6b software.

For 4-pulse-DEER, the samples were exchanged into deuterated buffer containing 30% d8- glycerol, checked by cwESR and plunge frozen into liq. N_2_. The pulse ESR measurements were carried out at Q-band (∼34 GHz) on a Bruker E580 spectrometer equipped with a 10 W solid state amplifier (150W equivalent TWTA) and an arbitrary waveform generator (AWG). DEER was carried out using four pulses (π/2-τ1-π-τ1-πpump-τ2-π-τ2-echo) with 16-step phase cycling at 60 K. The pump and probe pulses were separated by 56 MHz (∼20 G). The DEER data was background subtracted with an exponential background and the distance reconstruction was carried out using denoising and the SVD method (Srivastava, Madhur, et al., 2017; Srivastava, M., et al., 2017). Uncertainty in the distance distributions were calculated using the method described by Srivastava and Freed (Srivastava, M., et al., 2019). Briefly, the uncertainty we are plotting is showing the “range” of singular values over which the singular value decomposition (SVD) solution remains converged. For most of the data displayed in this paper we only used the first few singular values (SVs) and the solution remained converged for ± 1 or 2 SVs near the optimum solution. For example, if the optimum solution was 4 SVs then the range in which the solution remained converged is ∼3-6 SVs. We plot three lines - lowest range of SVs, highest range of SVs and optimum number of SVs – in the SI figures the optimum SV solution is shown in black and the region between the converged solutions with the highest and lowest number of SVs is shaded in red. Owing to the point-wise reconstruction of the distance distribution, the SVD method enables localized uncertainty at each distance value. Therefore, some points will have high uncertainty, whereas others low.

#### Turbidity Assays

Cysteine-less (Cys-less) FRQ was created in order to minimize disulfide cross-linking at the high protein concentrations needed for *in vitro* phase separation experiments. For this, all four of the native cysteine residues were substituted to serine using standard molecular biology techniques. For the phase separation experiments, this cys-less FRQ variant was purified and concentrated for dialysis into phase separation buffer. Cys-less FRQ was dialyzed overnight into 25 mM Tris pH 8.0, 150 mM NaCl or 25 mM Tris pH 8.0, 500 mM NaCl for p-FRQ and np-FRQ, respectively, at 4 ^°^C. Phase separation was induced by dialyzing in either low or high salt and increasing temperature. The protein solution can be observed as a clear solution before dialysis and then turbid after dialysis indicating LLPS has occurred. Microscopy (See *DIC and Fluorescence Microscopy*) confirmed the turbid solution is due to LLPS and not aggregation. Turbidity as measured by absorbance at 300 - 600 nm was then recorded by an Agilent Cary 3500 UV-Vis

Multicell Peltier using a temperature ramp rate of 5 °C/min increasing from 10 °C to 80 °C. A final protein concentration of 5-15 μM was used for all turbidity assays.

#### DIC and Fluorescence Microscopy

Phase separation was induced via overnight dialysis into 25 mM Tris pH 8.0, 150 mM NaCl or 25 mM Tris (pH 8.0), 500 mM NaCl for p-FRQ and np-FRQ, respectively. The final protein concentration used for microscopy were 5-15 μM. Images were taken on the Zeiss Axio Imager Z1 Upright Trinocular Fluorescence Microscope at 100x magnification. To test the interaction of FRQ with FRH and CK1, the phase separated FRQ was spiked with equimolar Cy5-labeled FRH or Cy5-labeled CK1 for final concentrations of 5 μM.

#### In vitro Phosphorylation Assay

22 μL samples containing 10-15 μM LFRQ were incubated in the presence or absence of 2 μM CK1 and 2 μM FRH in 50 mM Na_2_HPO_4_ pH 7.5, 300 mM NaCl, 200 mM arginine, 10 mM MgCl_2_. For the LLPS assay, FRQ was dialyzed into a buffer containing 50 mM Na_2_HPO_4_ pH 7.5, 300 mM NaCl and 10 mM MgCl_2_. 2 μL of 2.5 mM cold ATP and 0.1 µM of [γ-32P] ATP (3000 Ci/mmol, PerkinElmer) was added to produce a total volume of ∼25 μL. After ∼1h of incubation, the reaction was quenched with 25 μL of 4x SDS with 10 mM EDTA pH 8.0 and then subjected to gel electrophoresis on a 4–20% gradient Tris-glycine gel. The gel was dried with a GelAir dryer (Bio-Rad) and placed in an imaging cassette for at least 16 h and then imaged with a phosphorimager (GE, Typhoon FLA 7000). Band intensities we quantified using the Fiji software.

#### Mass Spectrometry and Phospho-Peptide Analysis

Phosphorylated FRQ was run on a 4-20% tris-glycine SDS-PAGE gel and the corresponding band was excised and trypsin digested. The digests were reconstituted in 0.5% formic acid (FA) for nanoLC-ESI-MS/MS analysis. The analysis was carried out using an Orbitrap Fusion^TM^ Tribrid^TM^ (Thermo-Fisher Scientific, San Jose, CA) mass spectrometer equipped with a nanospray Flex Ion Source, and coupled with a Dionex UltiMate 3000 RSLCnano system (Thermo, Sunnyvale, CA). The peptide samples (5 μL) were injected onto a PepMap C-18 RP nano trapping column (5 µm, 100 µm i.d x 20 mm) and then separated on a PepMap C-18 RP nano column (2 µm, 75 µm x 25 cm) at 35 °C. The Orbitrap Fusion was operated in positive ion mode with spray voltage set at 1.5 kV and source temperature at 275 °C. External calibration for FT, IT and quadrupole mass analyzers was performed. In data-dependent acquisition (DDA) analysis, the instrument was operated using FT mass analyzer in MS scan to select precursor ions followed by 3 second “Top Speed” data-dependent CID ion trap MS/MS scans at 1.6 m/z quadrupole isolation for precursor peptides with multiple charged ions above a threshold ion count of 10,000 and normalized collision energy of 30%. MS survey scans at a resolving power of 120,000 (fwhm at m/z 200), for the mass range of m/z 375-1600. Dynamic exclusion parameters were set at 35 s of exclusion duration with ±10 ppm exclusion mass width. All data were acquired under Xcalibur 4.4 operation software (Thermo-Fisher Scientific). The DDA raw files with MS and MS/MS were subjected to database searches using Proteome Discoverer (PD) 2.4 software (Thermo Fisher Scientific, Bremen, Germany) with the Sequest HT algorithm. The database search was conducted against a *Neurospora Crassa* database and *E. coli* database with adding LFRQ sequence into the database. The peptide precursor tolerance was set to 10 ppm and fragment ion tolerance was set to 0.6 Da with 2 miss-cleavage allowed for either trypsin or chymotrypsin. Variable modification of methionine oxidation, deamidation of asparagines/glutamine, phosphorylation of serine/threonine/tyrosine, acetylation, M-loss and M-loss+acetylation on protein N-terminus and fixed modification of cysteine carbamidomethylation, were set for the database search. Only high confidence peptides defined by Sequest HT with a 1% FDR by Percolator were considered for confident peptide identification. All the mass spectra of phosphorylated peptides were double checked by manually inspection.

#### Integrated Bioinformatic Analyses and Computational Biology

All canonical protein and UniRef50 sequences for FRQ, FRH and CK1, and SWISS-Prot obtained proteome sequences of *Neurospora crassa*, *Drosophila melanogaster* and *Homo sapiens* were downloaded from the UniProt database (Bateman, et al., 2021) A combination of **(a)** in-house written Python scripts to allow us to analyze large datasets, **(b)** localCIDER (Holehouse, et al., 2017), **(c)** IUPred2A (Mészáros, et al., 2018), **(d)** Agadir (Muñoz, et al., 1994), **(e)** AlphaFold 2.0 (Jumper, et al., 2021), **(f)** PScore (Vernon, et al., 2018) and (g) catGRANULE (Bolognesi, et al., 2016) were used to perform integrated bioinformatics analyses and computational biological studies of our proteins of interest. Unless mentioned, all programs were used by applying the default parameters. All scripts used to generate the data in this manuscript are provided as SI Datasets.

#### Neurospora crassa Strain Generation

Construction of the *frq[mNeonGreen]*, *son-1[mApple]* strain used for imaging was previously described (Bartholomai, 2022). Briefly, a plasmid was constructed to target the endogenous *frq* locus with DNA encoding *N. crassa* codon-optimized mNeonGreen appended to the C-terminus of the *frq* ORF with a flexible linker. A gene encoding hygromycin B phosphotransferase (HPH) was introduced downstream from the 3’ UTR of *frq* for selection. A plasmid was constructed containing the *N. crassa son-1* ORF with mApple appended to the C-terminus with a flexible linker for insertion at the *csr-1* locus. Transformation cassettes were amplified by PCR using NEB Q5® High-Fidelity 2X Master Mix (New England Bio Labs Cat. # M0492) and integrated into the genome by homologous recombination after electroporation of conidiophores. A homokaryotic strain containing the fluorescent fusion proteins was used for imaging.

#### Neurospora crassa Growth

*N. crassa* homokaryons containing *frq[mNeonGreen]* and *csr-1::son-1[mApple]*, were grown overnight on 2% agar gel pads made with Vogel’s minimal medium (Vogel, 1956) in constant light at 25 °C. Before imaging, a section near the edge of the growing mycelium was removed and inverted on a chambered cover glass (Thermo Scientific, Cat. # 155360).

#### Live Cell Image Acquisition and Processing

Images were acquired using a Zeiss AxioObserver laser scanning confocal microscope with a 100X/1.46 NA Oil immersion Plan-Apochromat objective. 561 nm and 488 nm lasers were used at 2% power with 2X averaging and a gain of 800 for detection of fluorescent signals by GaAsp PMTs. Images were processed for display using FIJI (Schindelin, et al., 2012). The cropped images displaying a single nucleus were enlarged 10X using bicubic interpolation and contrasted to the minimum and maximum gray level values. A multi-color lookup table was applied to the FRQ[mNeonGreen] channel to better observe heterogeneity of the signal, and the SON- 1[mApple] channel was pseudo-colored cyan to better contrast with the lookup table applied to the FRQ[mNeonGreen] channel. The surface plot of FRQ[mNeonGreen] was generated from the raw image. Movie S1, and the image in Figure 6E were acquired using a Nikon Ti-E based system, equipped with an Andor W1 spinning disk scan head, an Andor Zyla sCMOS camera, a 488 nm laser, and Chroma bandpass filters. Figure 6E is a still image from a single focal plane from a z-stack that was deconvolved using Nikon Elements and was acquired using a 100X/NA1.45 plan apo oil immersion objective. Figure 6E and Movie S1 were acquired using a 60X/NA1.45 plan apo oil immersion objective. The 1-minute movie was acquired with a 300 ms exposure per frame. Image processing for display was carried out using FIJI.

### Results

**Table S1:** SAXS parameters for p-FRQ, np-FRQ and their reconstitution into the FFC.

| | $R_g$ [Guinier] (Å) | $R_g$ [P(r)] (Å) | Dmax (Å) | MW [$V_c$] (kDa) |
| --- | --- | --- | --- | --- |
| p-FRQ | $123 \pm 7$ | $142 \pm 5$ | $530 \pm 60$ | 816 |
| np-FRQ | $100 \pm 2$ | $116 \pm 4$ | $446 \pm 40$ | 680 |
| p-FFC | $101 \pm 2$ | $136 \pm 5$ | $510 \pm 20$ | 314 |
| np-FFC | $80 \pm 2$ | $109 \pm 7$ | $480 \pm 10$ | 286 |

MW calculations for p-FRQ and np-FRQ are inaccurate because $V_c$ (Volume of Correlation) plots (Integrated area of $I(q)q$ vs $q$ ; $q = 2\pi/d$ ) did not converge for these species. Expected MW for LFRQ is 110 kDa. Expected FFC MW for the 1:1:1 stoichiometry of FRQ:FRH:CK1 is 273 kDa. Errors in Dmax were estimated from the range for which $P(r)$ tailed to zero in the high distance range without constraint.

**Table S2:** FRQ Cysteine residue locations and context.

| Site | Domain | Binding Partner at this region | Reference |
| --- | --- | --- | --- |
| 370C | N/A |  |  |
| 490C | FCD2 | CK1 | (Querfurth, Christina, et al., 2011) |
| 658C | N/A |  |  |
| 746C | FFD | FRH | (Cheng, et al., 2005) |

**Table S3:** FRQ regions that bind clock proteins.

| Binding Partner | Region | Cysteine residue(s) present in this region | Reference |
| --- | --- | --- | --- |
| FRH | 695-778 | 746C | (Cheng, et al., 2005) |
| CK1 | 319-326; 488-495 | 490C | (Querfurth, Christina, et al., 2011) |
| WC2 |  | 490C, 658C, 746C | (Guo, et al., 2010) |

**Table S4:** δ values from the cw-ESR central peak for FRQ variants of Fig. 3C.

| Site | np-FRQ (G) | p-FRQ (G) |
| --- | --- | --- |
| NT | 1.75 | 1.87 |
| 490C | 1.97 | 2.47 |
| 658C | 1.97 | 2.01 |
| 746C | 1.76 | 1.82 |
| CT | 2.30 | 1.70 |

**Table S5:**
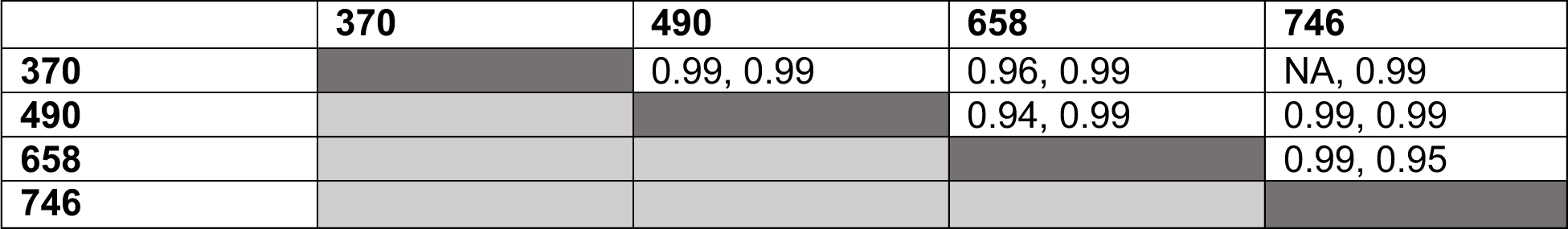
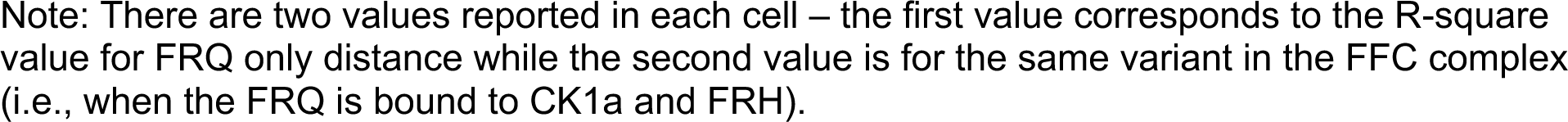
R-square values for the quality of the two component gaussian fits to the distance distributions.

|  | 370 | 490 | 658 | 746 |
| --- | --- | --- | --- | --- |
| 370 |  | 0.99, 0.99 | 0.96, 0.99 | NA, 0.99 |
| 490 |  |  | 0.94, 0.99 | 0.99, 0.99 |
| 658 |  |  |  | 0.99, 0.95 |
| 746 |  |  |  |  |

Note: There are two values reported in each cell – the first value corresponds to the R-square value for FRQ only distance while the second value is for the same variant in the FFC complex (i.e., when the FRQ is bound to CK1a and FRH).

**Table S6:**
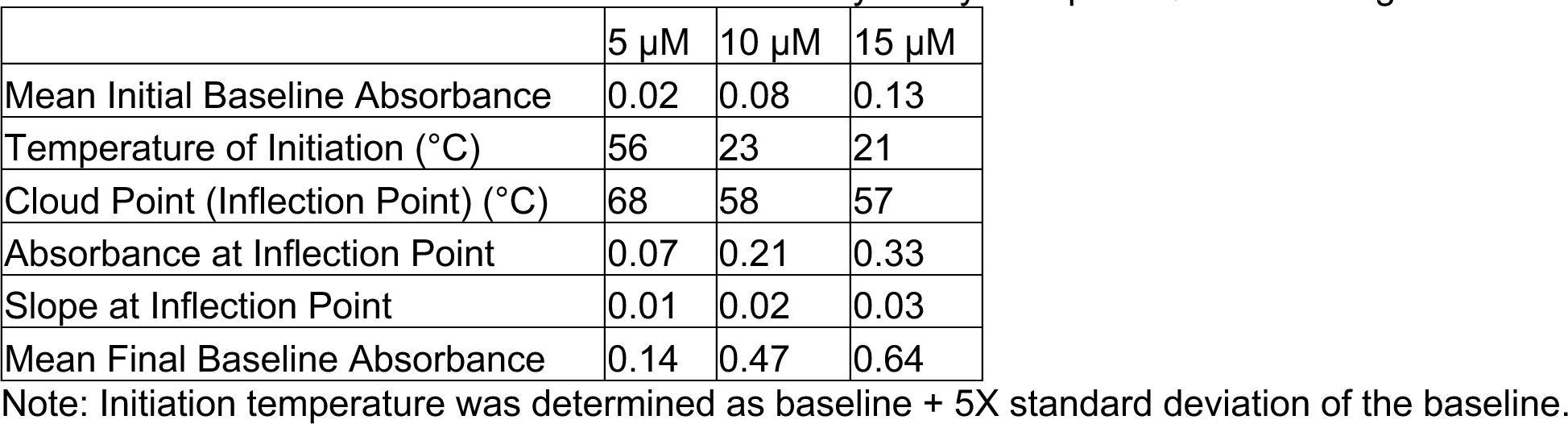
Calculated values from the UV-Vis turbidity assays of np-FRQ shown in Figure 6C.

| | 5 μM | 10 μM | 15 μM |
| --- | --- | --- | --- |
| Mean Initial Baseline Absorbance | 0.02 | 0.08 | 0.13 |
| Temperature of Initiation ($^{\circ}$C) | 56 | 23 | 21 |
| Cloud Point (Inflection Point) ($^{\circ}$C) | 68 | 58 | 57 |
| Absorbance at Inflection Point | 0.07 | 0.21 | 0.33 |
| Slope at Inflection Point | 0.01 | 0.02 | 0.03 |
| Mean Final Baseline Absorbance | 0.14 | 0.47 | 0.64 |

Note: Initiation temperature was determined as baseline + 5X standard deviation of the baseline.

**Table S7:** Calculated values from the UV-Vis turbidity assays of p-FRQ shown in Figure 6C.

| | 5 μM | 10 μM | 15 μM |
| --- | --- | --- | --- |
| Mean Initial Baseline Absorbance | 0.09 | 0.21 | 0.46 |
| Temperature of Initiation ($^{\circ}$C) | 45 | 42 | 24 |
| Cloud Point (Inflection Point) ($^{\circ}$C) | - | - | - |
| Absorbance at Inflection Point | - | - | - |
| Slope | 0.00 | 0.01 | 0.02 |
| Mean Final Baseline Absorbance | - | - | - |

Note: Initiation temperature was determined as baseline +5X standard deviation of the baseline. Also, because p-FRQ continues to undergo LLPS within the temperature range of the experiment, we cannot calculate a cloud point.

**Figure S1:**
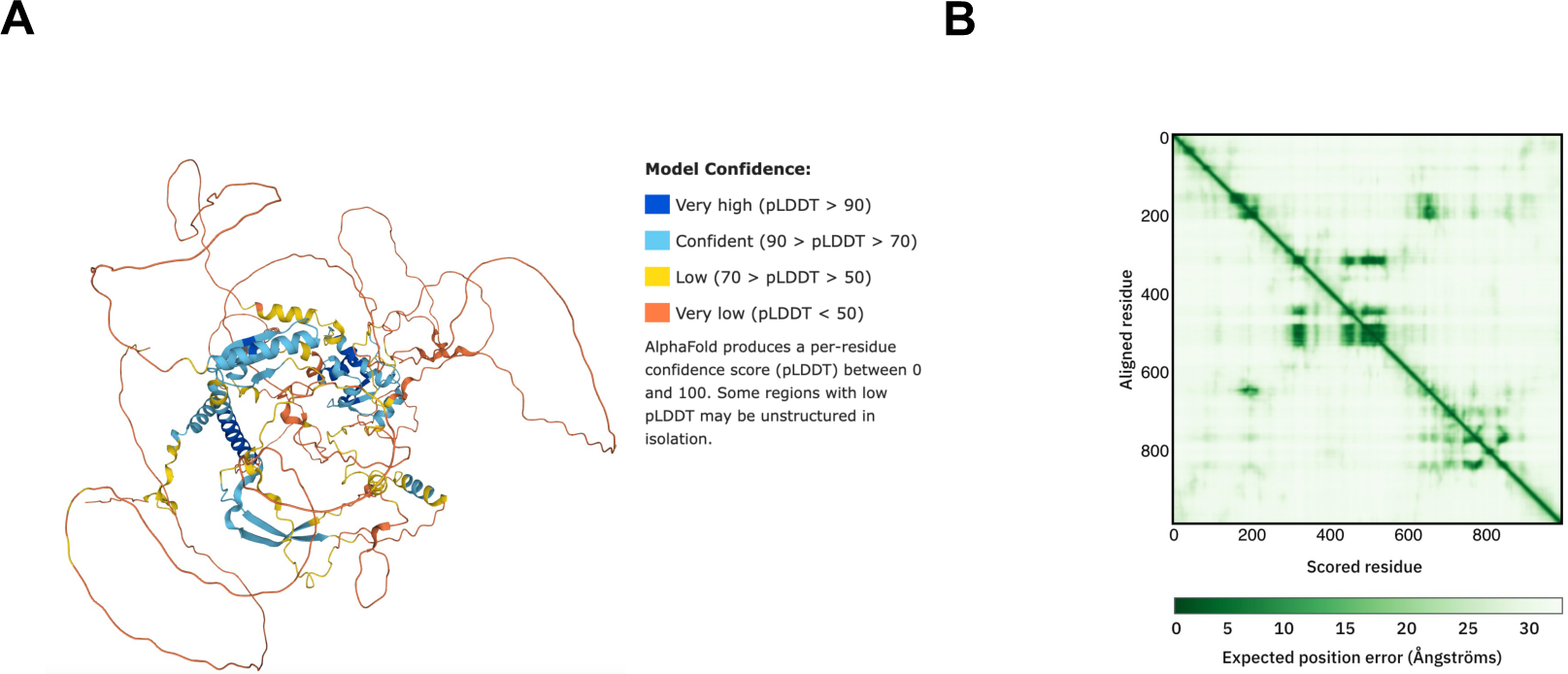
(A) AlphaFold model of LFRQ colored by pLDDT score. (B) Predicted error alignment plot that shows the uncertainty in the relative positions of two residues in the structure. (Image source: https://www.uniprot.org/uniprotkb/P19970/entry#structure)

**Figure S2:**
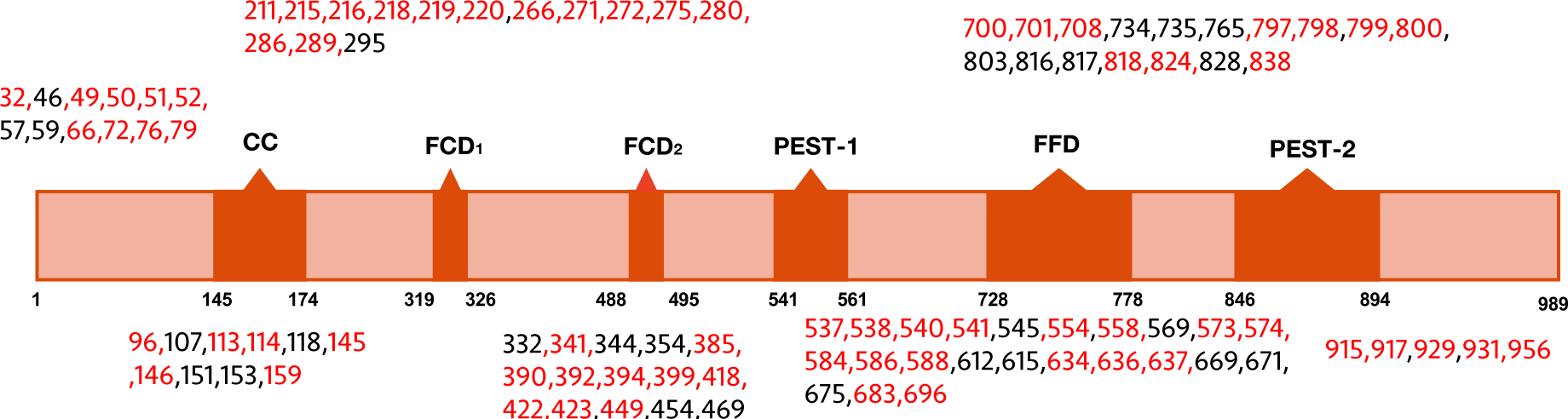
Domain map of FRQ with phospho-sites (as identified by mass-spec) highlighted. The sites in red were also identified in (Baker, et al., 2009; Tang, et al., 2009).

**Figure S3:**
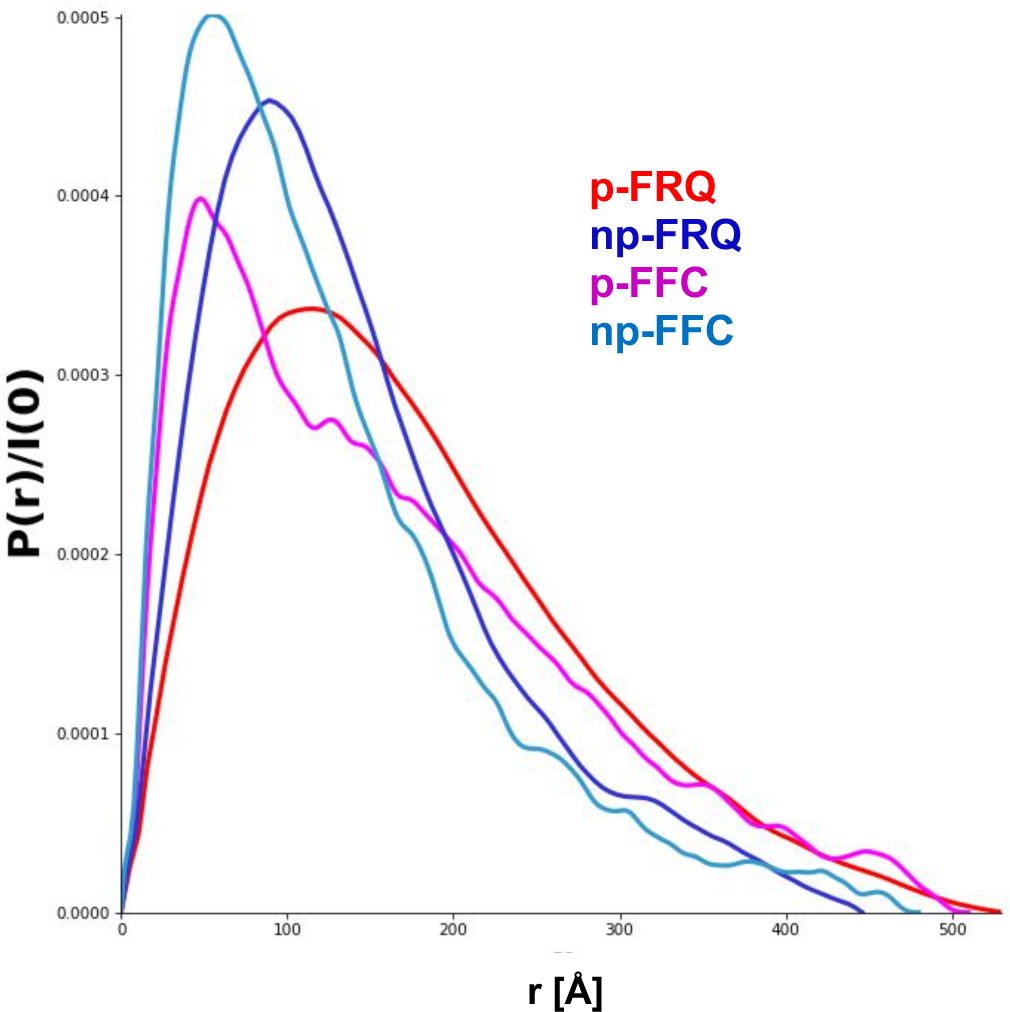
Pair-wise distance distributions calculated from SAXS data for p-FRQ (red), np-FRQ (dark blue), p-FFC (magenta), and np-FFC (light blue).

**Figure S4:**
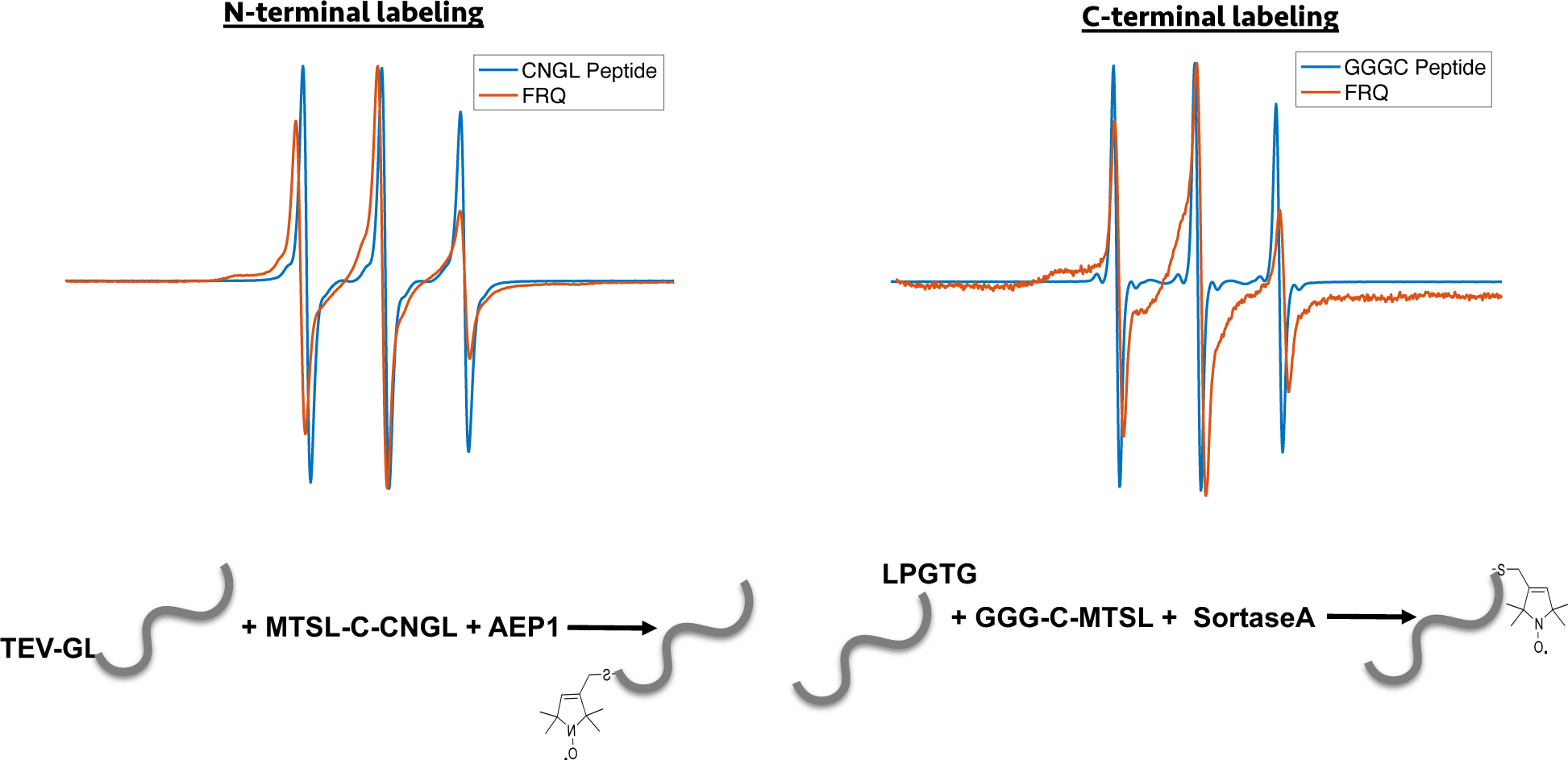
Enzymatic labeling of FRQ at its termini. X-band (9.8 GHz) RT cw-ESR measurements for the spin-labeled peptide by itself and when it has been grafted onto either the N or C-terminus of FRQ.

**Figure S5:**
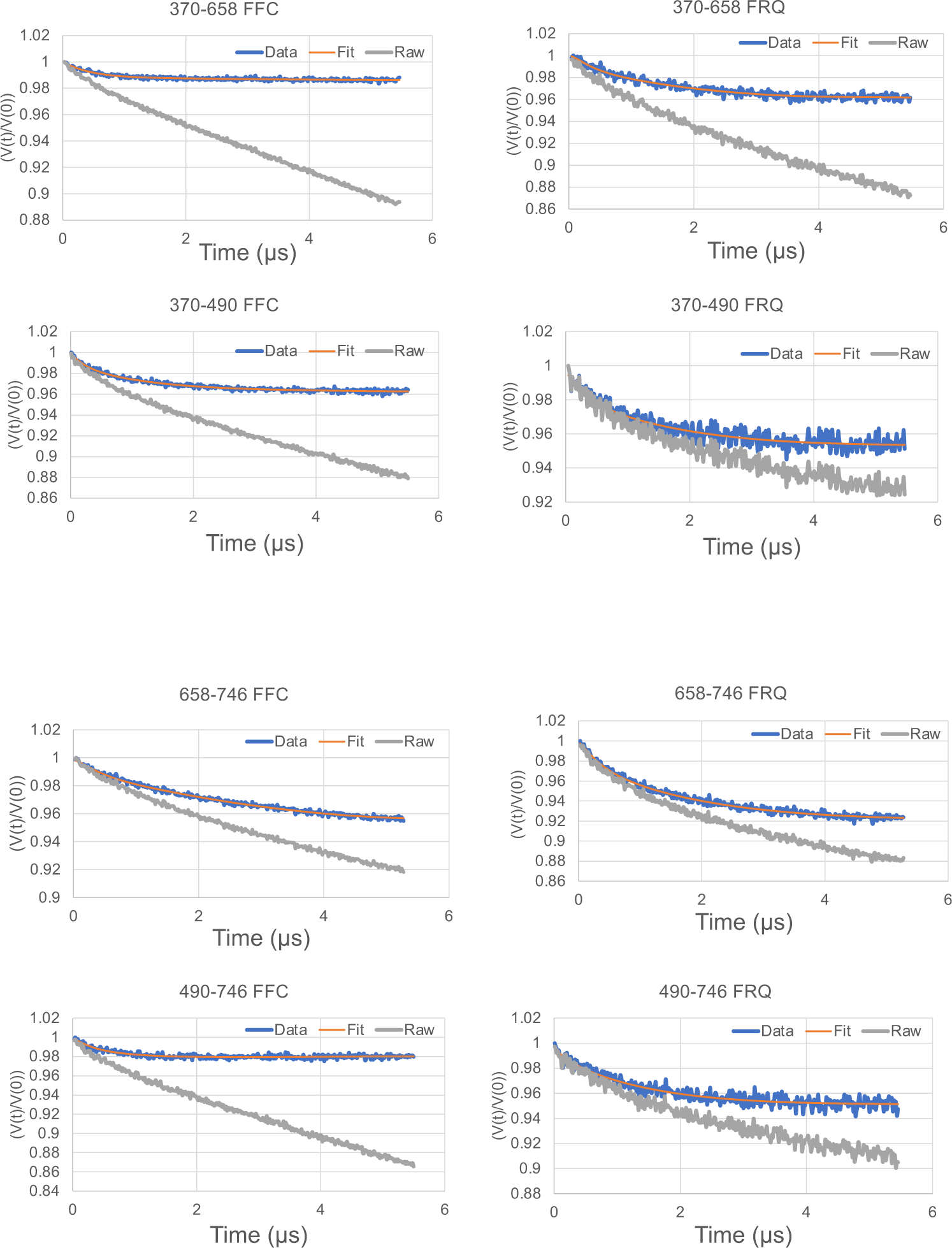
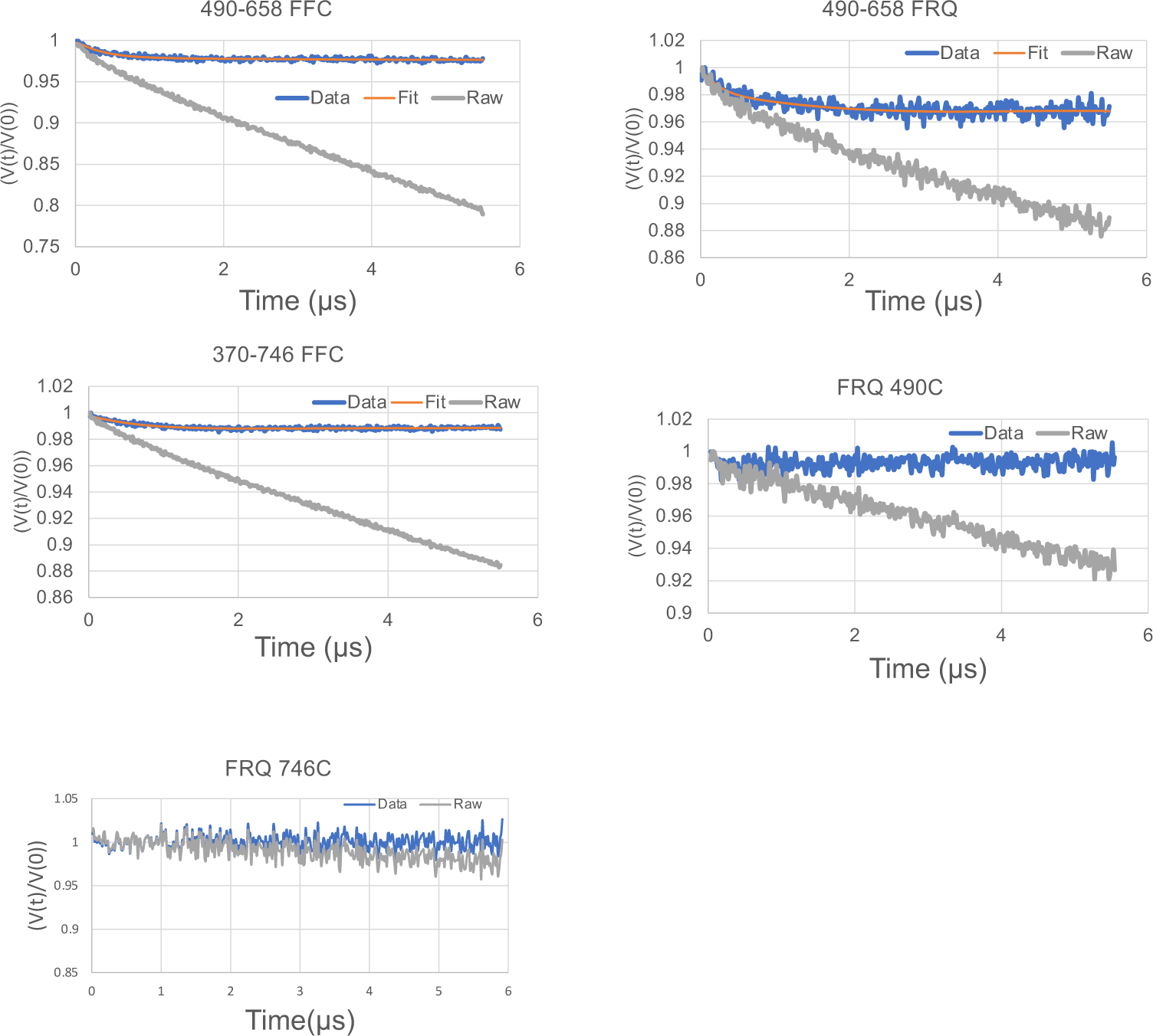
Time domain traces before (Raw) and after (Data) background correction and trace fits (Fit) from DEER experiments of different double-cysteine FRQ variants alone or within the FFC (i.e., when FRQ was bound to FRH and CK1a). The spins were present on the residue number of FRQ that is highlighted for example ‘370-490’ refers to FRQ spin-labeled at the 370C and 490C site and the DEER experiment probed the dipolar coupling of the spins at these sites. The last graphs show the time domain data for the control single cysteine DEER experiments with p-FRQ 490C and p- FRQ 746C.

**Figure S6:**
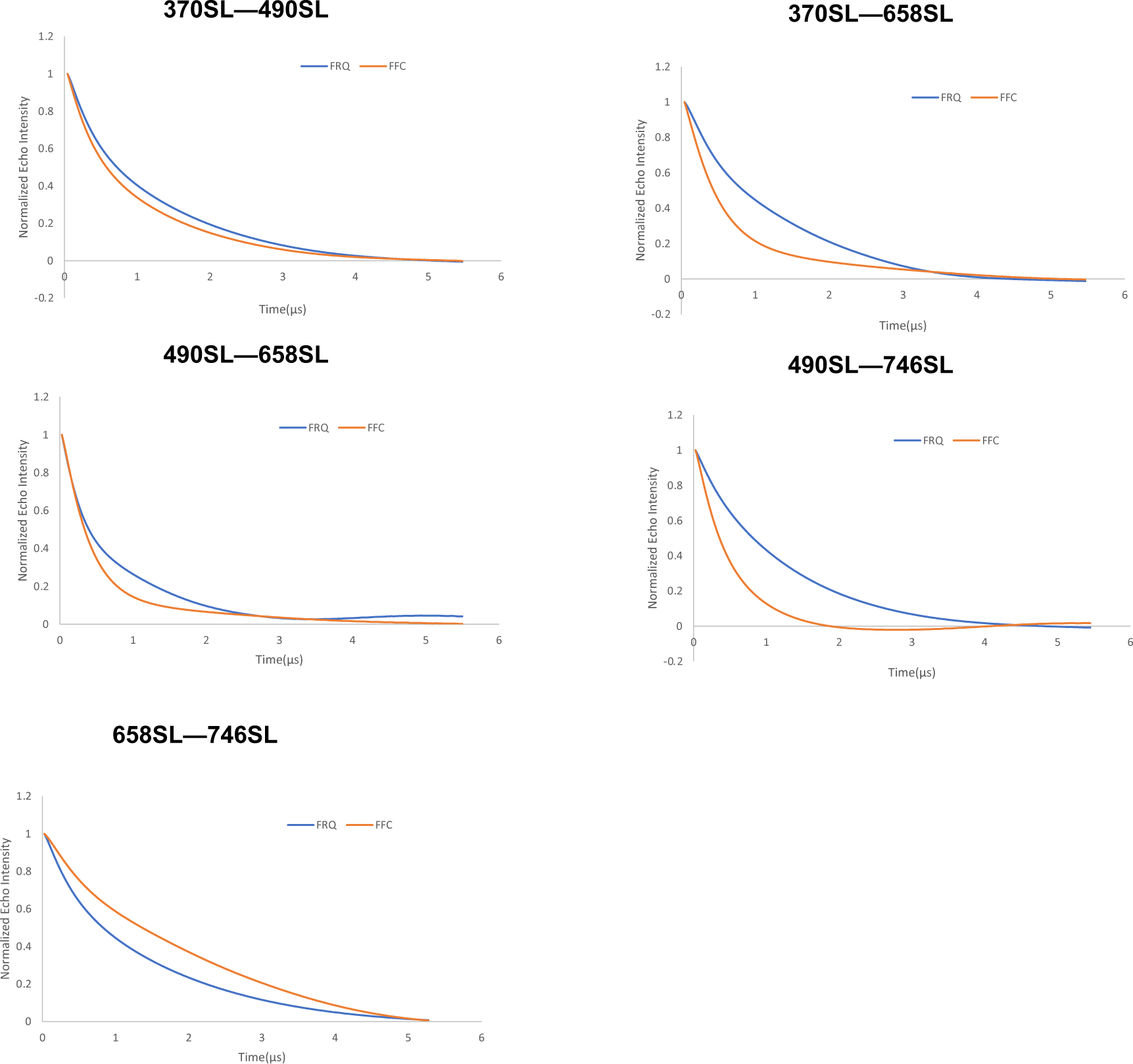
Baseline-corrected time domain fit traces from Q-band DEER experiments of different double-cysteine FRQ mutants alone (blue) and within the FFC (orange). Original data are shown in Supplementary Figures S5 and S9.

**Figure S7:**
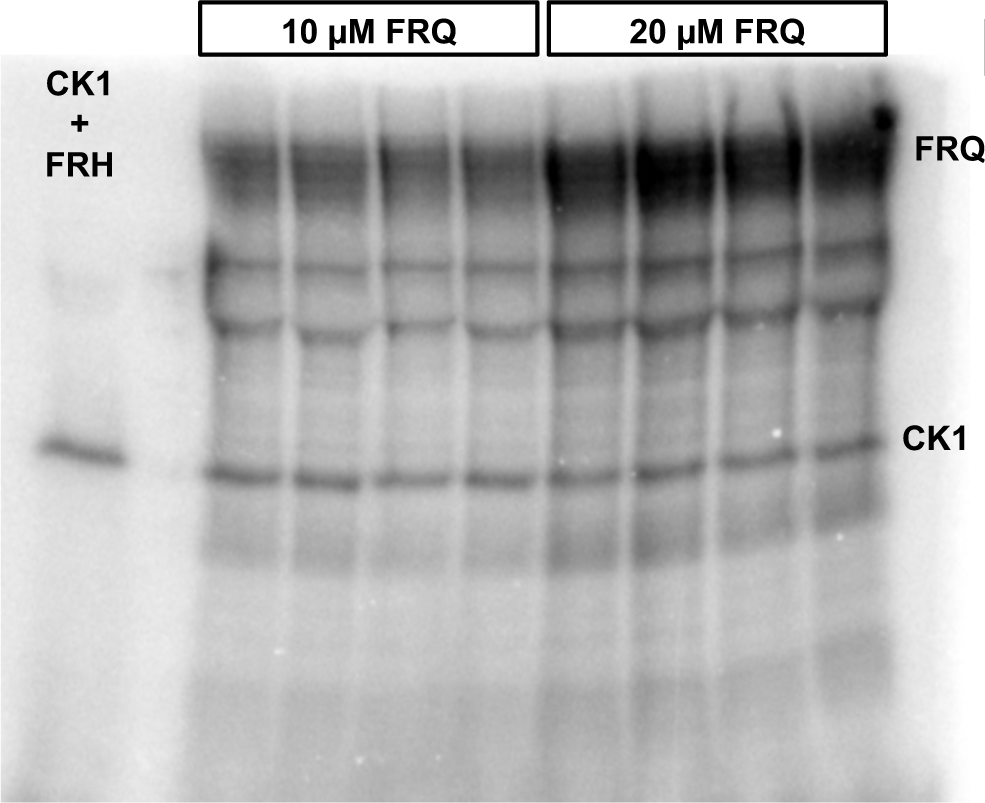
Full autoradiograph of np-FRQ phosphorylation by CK1 and γ-^32^P-ATP from Figure 4A inset. Four technical replicates are shown for each np-FRQ concentration.

**Figure S8:**
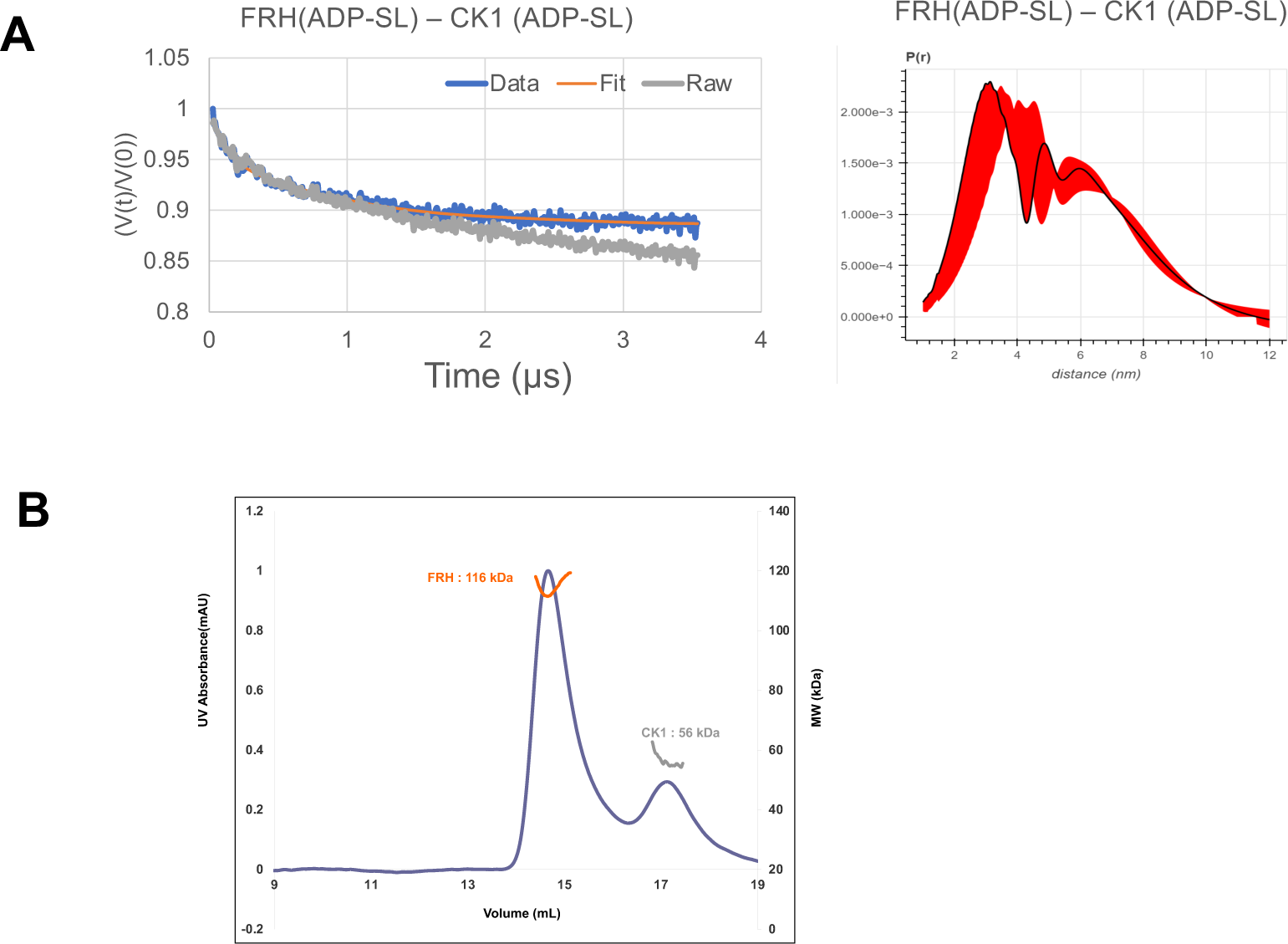
(A) Background-corrected time domain trace and error analyses from DEER experiments targeting the ATP-binding pockets of FRH and CK1. The DEER experiment probed the dipolar coupling of the spins in the ADP-β-S-SL molecule that was present in ATP-binding pocket of FRH or CK1. (B) Size-exclusion chromatography- multiangle light scattering (SEC-MALS) of a mixture of purified FRH and CK1. The proteins elute separately as monomers with no evidence of co-migration.

**Figure S9:**
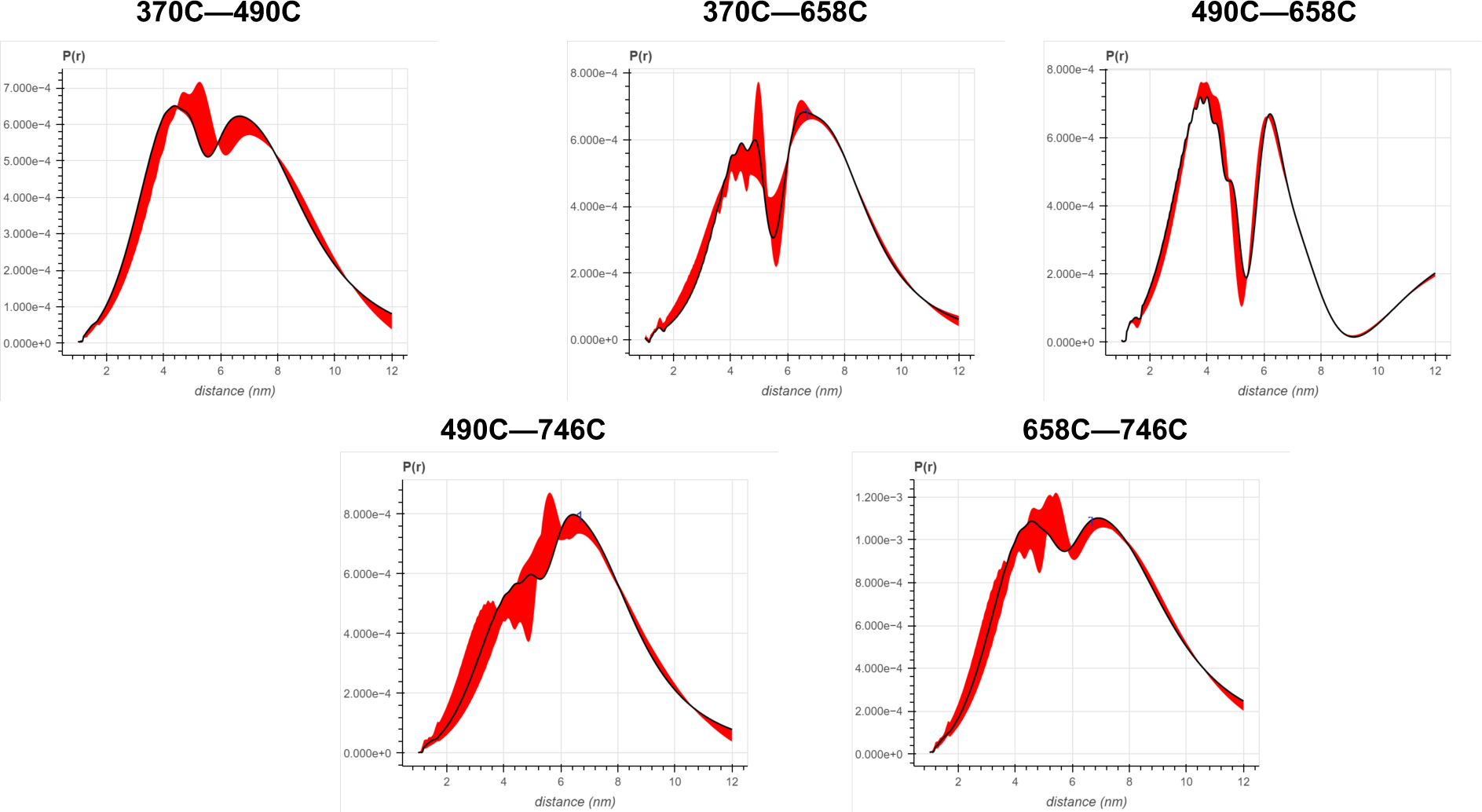
Error analysis of the distance distributions produced from DEER experiments of different double-cysteine FRQ variants. The spins were present on the amino acid number of FRQ that is highlighted for example ‘370-490’ refers to FRQ spin-labeled at the 370C and 490C site and the DEER experiment probed the dipolar coupling of the spins at these sites.

**Figure S10:**
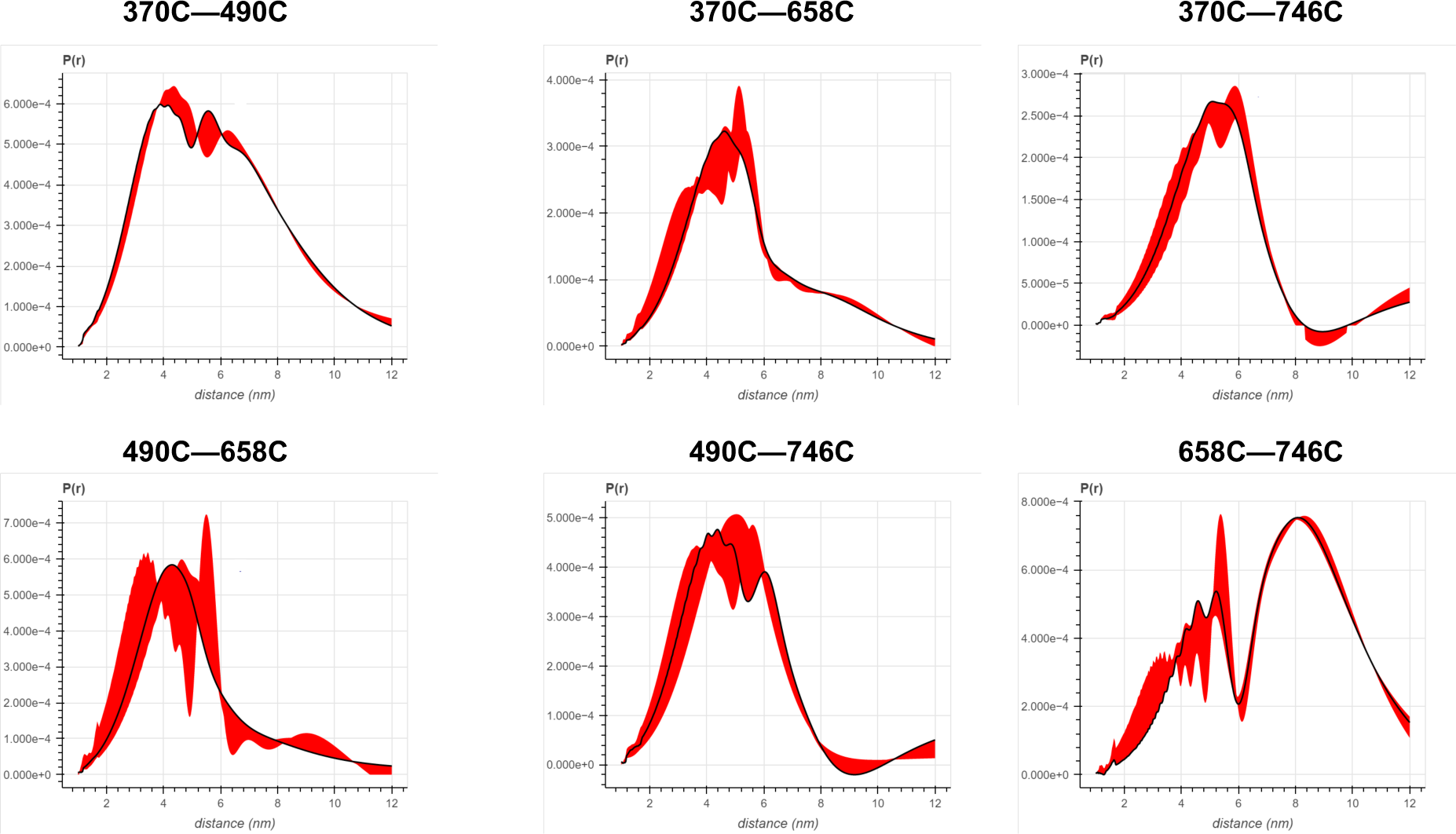
Error analysis of the distance distributions produced from DEER experiments of different double-cysteine FRQ variants within the FFC (i.e., when FRQ was bound to FRH and CK1). The spins were present on the amino acid number of FRQ that is highlighted for example ‘370-490’ refers to FRQ spin-labeled at the 370C and 490C site and the DEER experiment probed the dipolar coupling of the spins at these sites.

**Figure S11:**
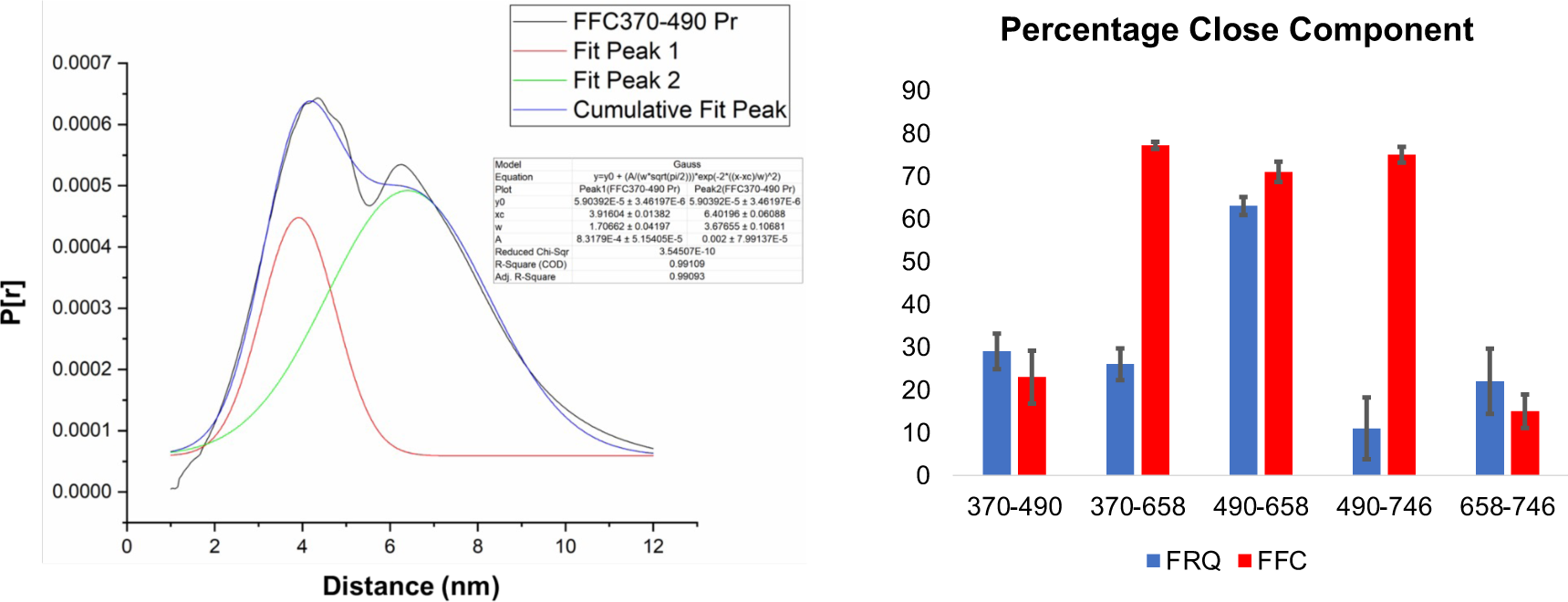
(Left)The two component Gaussian fitting of the distance distribution (P[r]) obtained from SVD for FFC 370-490 (i.e., spins were placed on resides 370 and 490 on FRQ, and this protein was bound to CK1 and FRH). No constraints were placed on the width, position, or amplitude of either component and the best fit values were obtained by minimizing the difference between the fits and the SVD distance distributions. Similar fits were carried out for all the different variants. The quality of the fit is reported by the R-square values and ranged between 0.94-0.99 for all the samples tested. They are reported in Table S5. (Right) The proportion of the closer distance peak in the distance distributions of FRQ (blue) or FRQ+FRH+CK1(FFC) (red) samples. The error bars reflect the uncertainty in estimating the population of each peak and were derived from uncertainties in the distance distributions shown in *SI Appendix*, Fig. S9 and S10.

**Figure S12:**
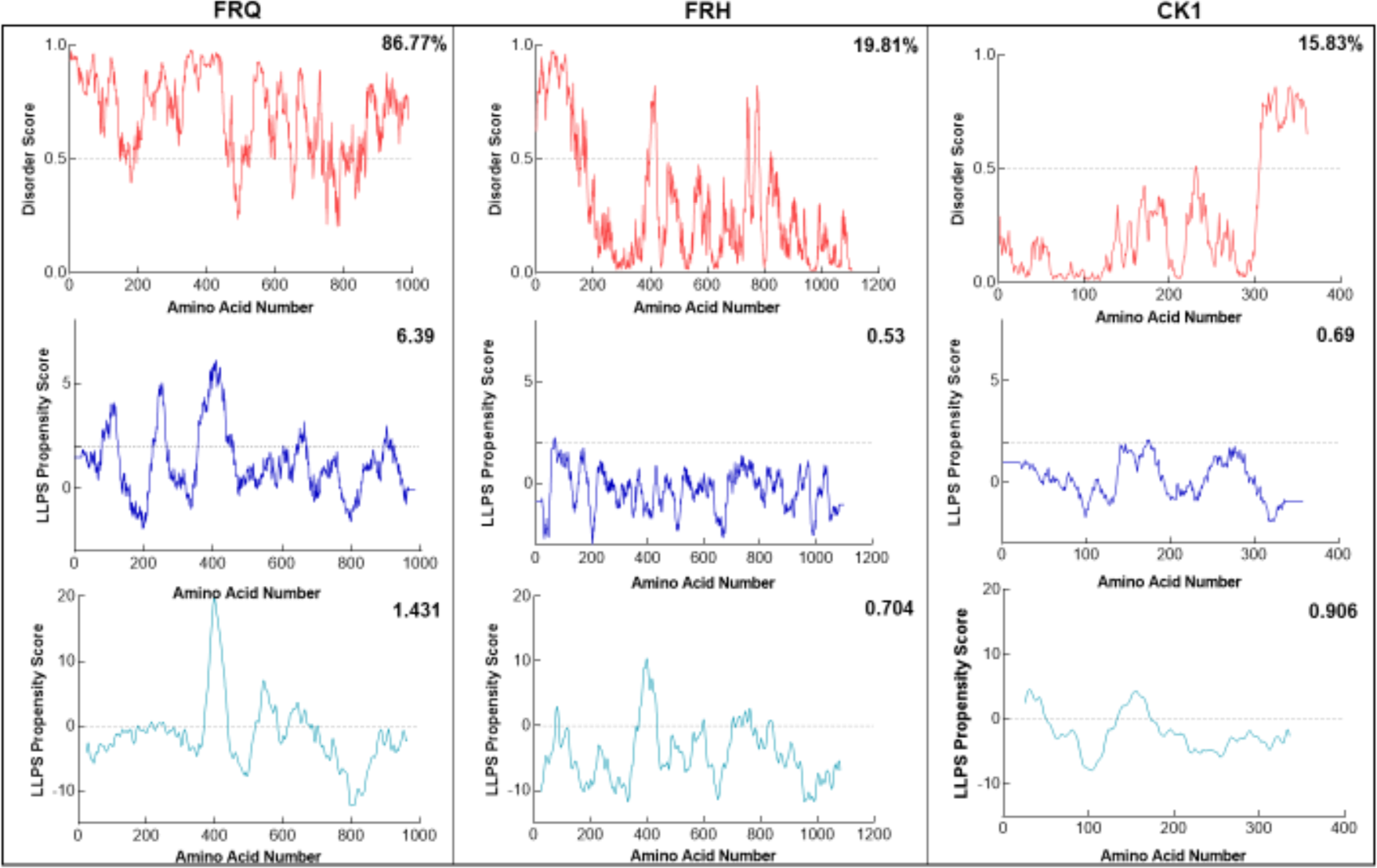
FRQ, but not FRH or CK1, is highly disordered and predicted to undergo LLPS. Shown are the disorder and LLPS propensity scores of the members for the FFC complex, FRQ, FRH and CK1. Columns from left to right are disorder and phase separation predictions for FRQ, FRH and CK1, respectively. Disorder graphs (red) were computed using the IUPred2A program (Mészáros, et al., 2018) where values above a 0.5 threshold (dashed line) are considered disordered. LLPS propensity predictions were computed using “Pi-Pi” (blue) (Vernon, et al., 2018) and catGRANULE (light blue) (Bolognesi, et al., 2016). Dashed lines represent neutral propensity. Shown on the top right corner of each graph are the percent disorder or LLPS propensity scores of each protein.

**Figure S13:**
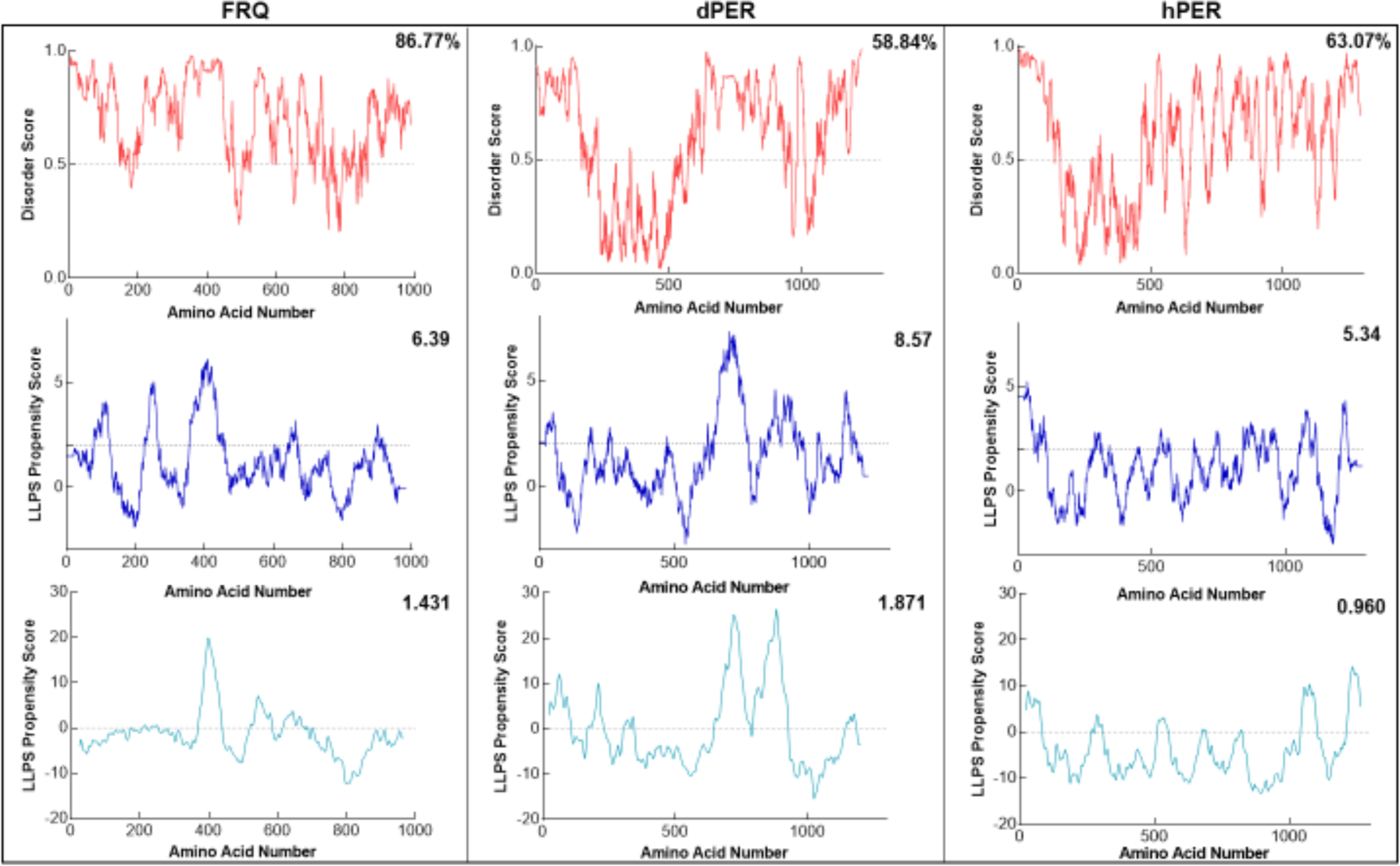
FRQ and its functional homologs are predicted to undergo LLPS. Shown are the disorder and LLPS propensity scores for FRQ, dPER1 and hPER1. Data for the disorder graphs (red) were computed using the IUPred2A program where values above a 0.5 threshold (dashed line) are considered disordered. Pi-pi predictions (blue) were computed using (Vernon, et al., 2018), while catGRANULE predictions (light blue) were computed using catGRANULE server (Bolognesi, et al., 2016). Shown on the top-right corner of each graph are the percent disorder, PScore or catGRANULE score for each protein. Dashed lines represent neutral propensity. FRQ, dPER1 and hPER1 are all predicted to be highly disordered (86.77%, 58.84% and 63.07%, respectively) and undergo LLPS.

**Figure S14:**
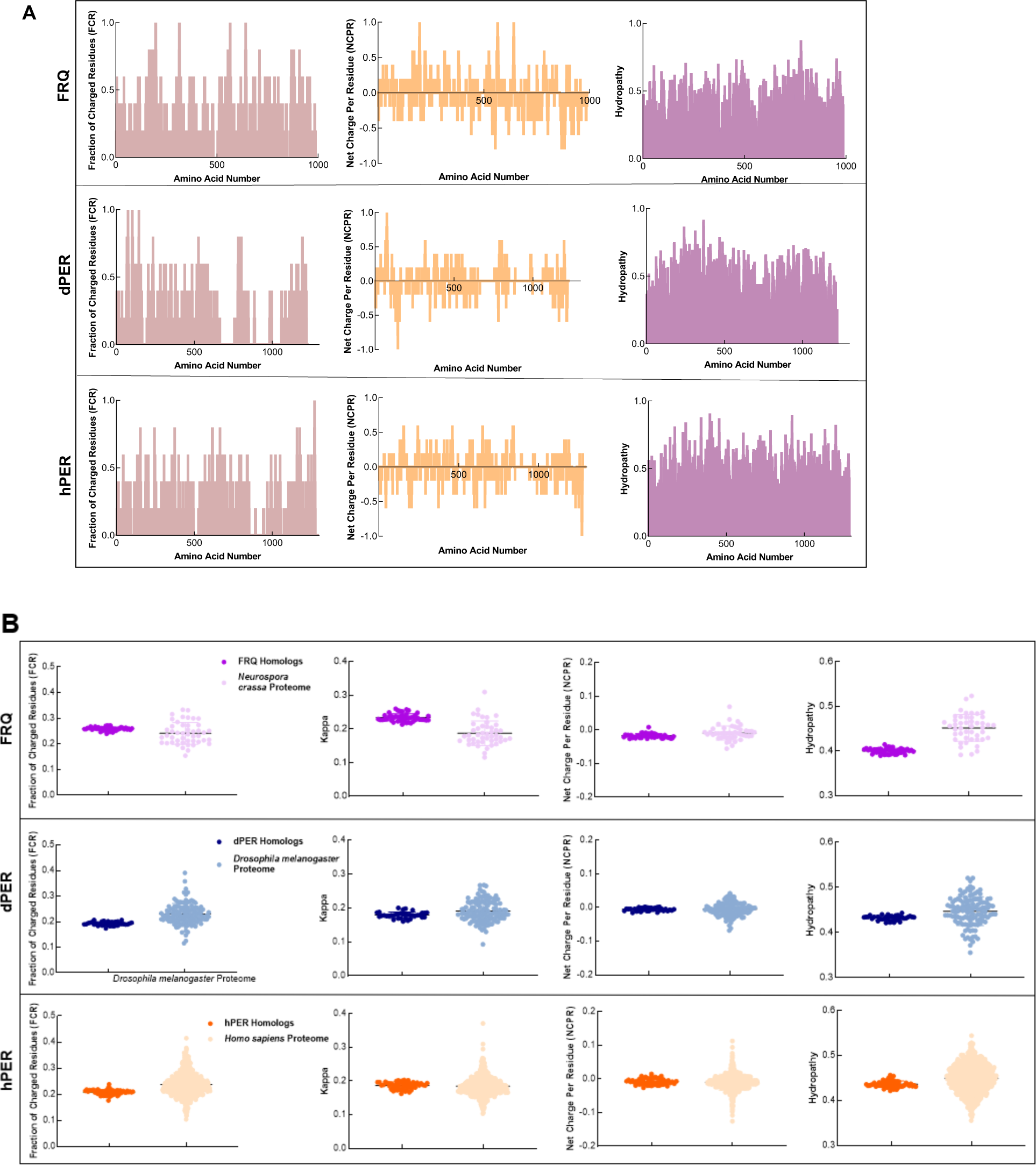
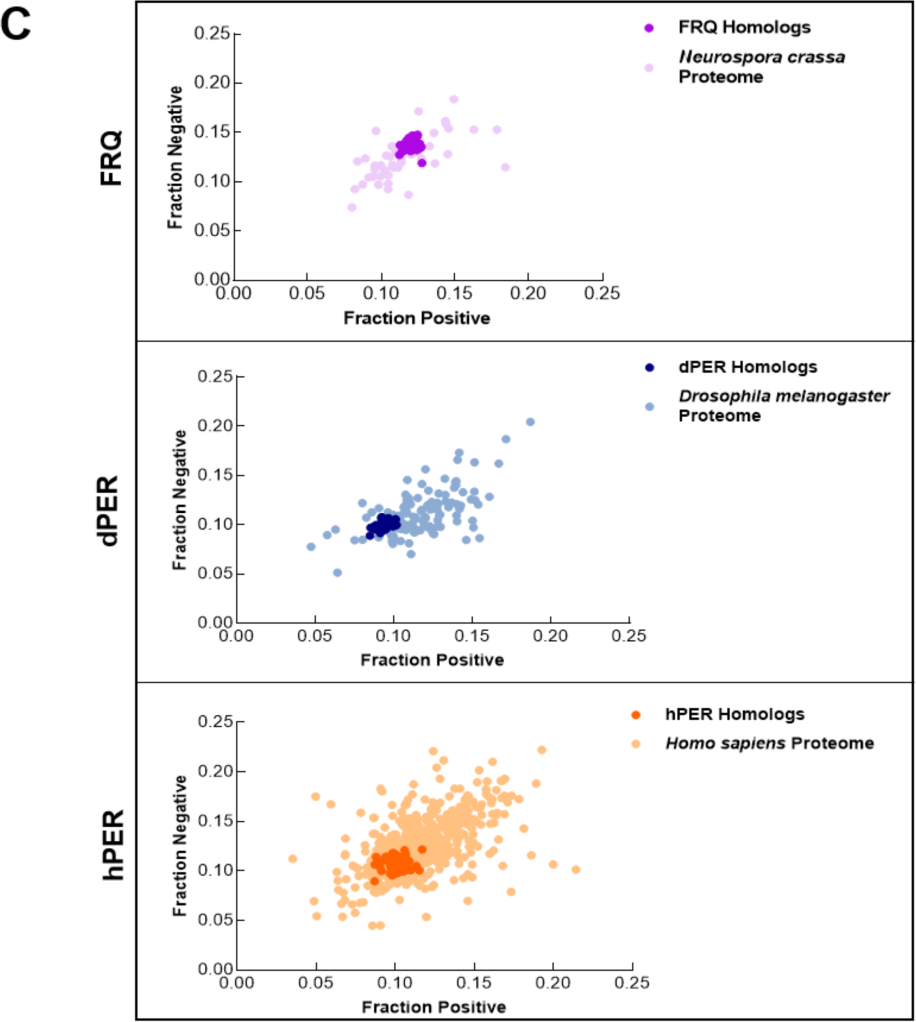
FRQ and its homologs, dPER1 and hPER1, have similarly disparate physicochemical properties compared to their proteomes. (A) Shown are the distribution of fraction of charged residues (FCR), net charge per residue (NCPR), and hydropathy along the sequence of FRQ, dPER1 and hPER1. (B) Sequence characteristics of FRQ and its homologs, dPER1 and hPER1. Various protein sequence parameters are calculated using localCIDER (Holehouse, et al., 2017) and displayed including fraction of charged residues (FCR), the degree of charge mixing (Kappa), net charge per residue (NCPR), and hydropathy (the hydrophobic and hydrophilic character of protein chain as defined by (Kyte, et al., 1982)) for the UniProtRef50 clusters of FRQ (purple), dPER1(blue) and hPER1 (orange). For comparison are shown protein sequences of similar length from the SWISS-Prot reviewed proteomes of *Neurospora crassa* (light purple), *Drosophila melanogaster* (light blue) and *Homo sapiens* (light orange). Hydropathy is consistently low for the clock proteins and FRQ orthologs have a high degree of charge separation, i.e., tracks of negative and positive residues (high kappa). High kappa values are common for proteins that phase separate (Somjee, et al., 2020). (C) Classification of the charge properties of the Ref50 clusters of FRQ and its functional homologs, dPER1 and hPER1. Das-Pappu phase plots showing the fraction of negatively vs positively charge residues for the UniProtRef50 clusters of FRQ (purple), *Neurospora crassa* proteome control (light purple), dPER1 (blue), *Drosophila melanogaster* proteome control (light blue), hPER1 (orange) and Homo sapiens proteome control (light orange). Shown are the fraction of negative residues versus fraction of positive residues for each cluster of homologs and their controls (similar size proteins from their respective proteomes). Despite the lack of amino acid conservation between the functional homologs of FRQ, dPER1 and hPER1, the ratio of fraction negative to fraction positive residues are similar among the clock proteins. dPER and hPER1 have a similarly low value of both fraction positive and fraction negative compared to their respective proteomes, whereas the fraction positive and fraction negative residues are both higher in FRQ. Note: Ref50 is a clustering method utilized by UniProt and is defined in (Steinegger, et al., 2018). While this work was in preparation, conceptually similar analyses were carried out by Jankowski et al., 2022 (Jankowski, et al., 2022).

**Figure S15:**
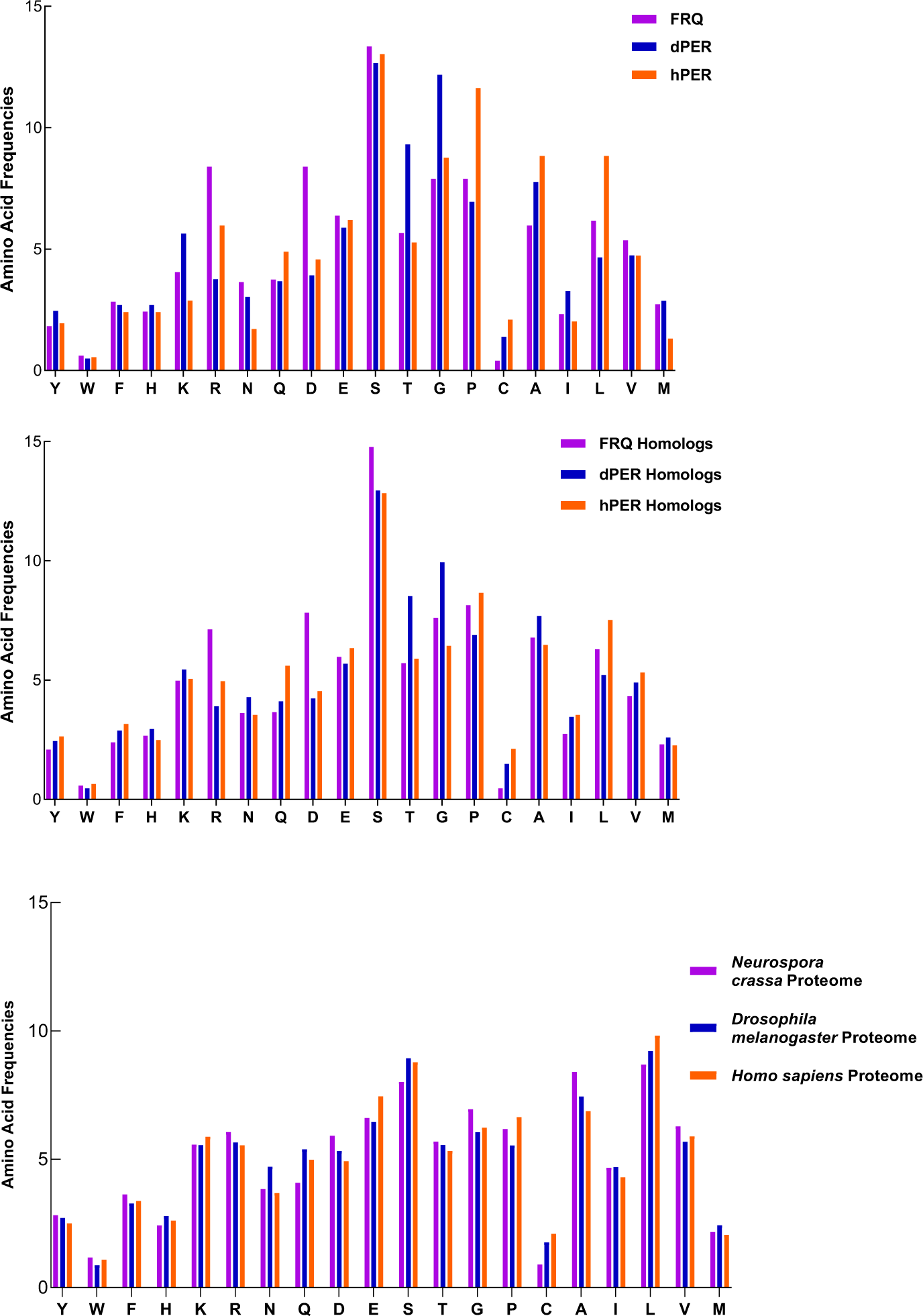
FRQ and its functional homologs dPER1 and hPER1 have similar amino acid composition compared to their respective proteomes. FRQ and its functional homologs dPER1 and hPER1 have similar amino acid compositions, which differs substantially from that of their respective proteomes. The amino acid composition for FRQ, dPER1 and hPER1 and their respective homolog clusters and proteomes are shown. Generally, G, S, T and P residues are enriched whereas hydrophobic (A, I, L, V) residues are relatively depleted for FRQ, dPER1, hPER1 and their homologs compared to their proteomes. Unlike dPER1 and hPER1 and their homologs, FRQ and its homologs are also highly enriched in R and D.

**Figure S16:**
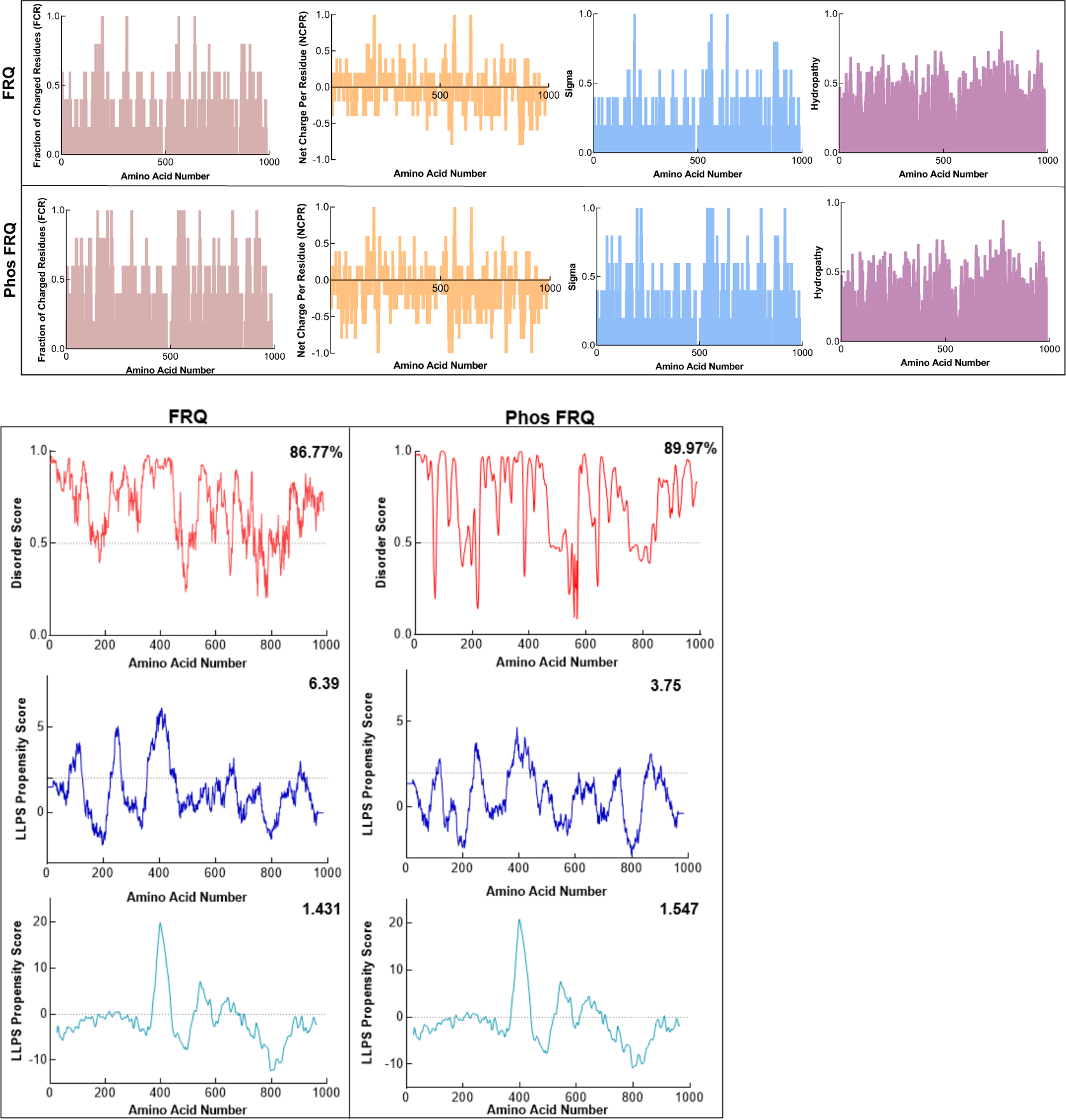
Phosphorylation modulates the physicochemical properties of FRQ. Shown are the distribution of fraction of charged residues (FCR), net charge per residue (NCPR), and hydropathy along the sequence of np and p-FRQ (top panel). In the bottom panel, shown are the disorder and LLPS propensity scores for np and p-FRQ. Data for the disorder graphs (red) were computed using the IUPred2A program (Mészáros, et al., 2018) where values above a 0.5 threshold (dashed line) are considered disordered. Pi-pi predictions (blue) were computed using, while catGRANULE predictions (light blue) were computed using catGRANULE server (Bolognesi, et al., 2016). Shown on the top-right corner of each graph are the percent disorder, PScore or catGRANULE score for each protein. Dashed lines represent neutral propensity. Note: Glu(E) residues were employed as phospho-mimetics in all analysis.

**Figure S17:**
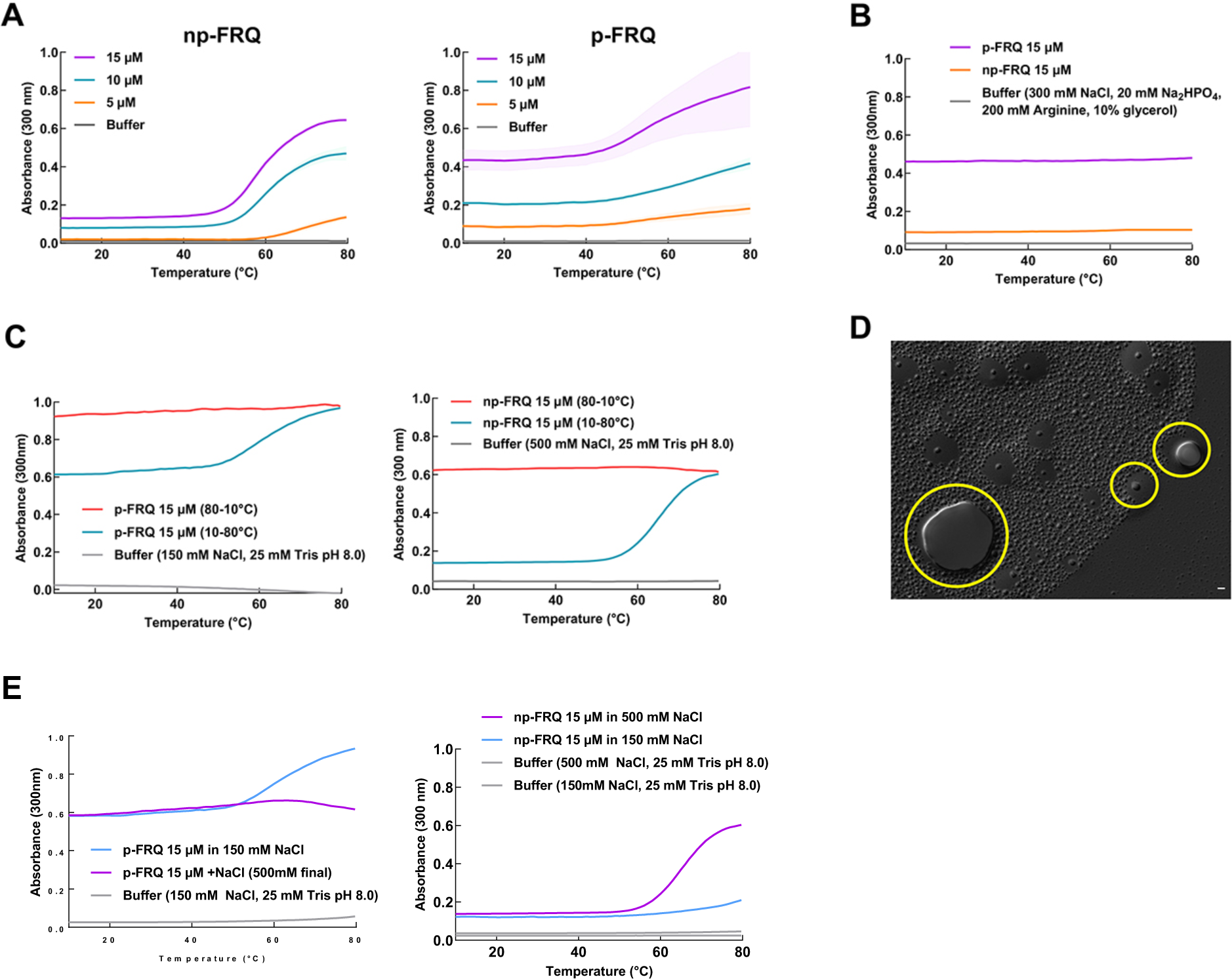
Exploring the phase behavior of FRQ. (A) UV-Vis absorption spectrum of p-FRQ (in 150 mM NaCl, 25 mM Tris pH 8.0) and np-FRQ (in 500 mM NaCl, 25 mM Tris pH 8.0) as a function of temperature at increasing concentrations. Note: the shading around each curve represents the standard deviation from the mean absorbance from 2 technical replicates. 15 μM p-FRQ had the greatest standard deviation as seen by the largest shaded area. (B) UV-Vis absorption spectrum of p-FRQ and np-FRQ in a non LLPS promoting buffer as a function of temperature. Note: p-FRQ appears to undergo a small amount of LLPS even in non-LLPS buffer as seen by its relatively high absorbance relative to np-FRQ.(C) (Left) UV-Vis absorption spectrum of p-FRQ (in 150 mM NaCl, 25 mM Tris pH 8.0) and (Right) np-FRQ (in 500 mM NaCl, 25 mM Tris pH 8.0) as a function of temperature to check for reversibility of LLPS transition. (D) DIC micrograph of FRQ showing droplets fusing and docking as indicated within the yellow circles. (E) UV-Vis absorption spectrum of p-FRQ (Left) and np-FRQ (Right) as a function of temperature in either low salt (150 mM) or high salt (500 mM).

**Figure S18:**
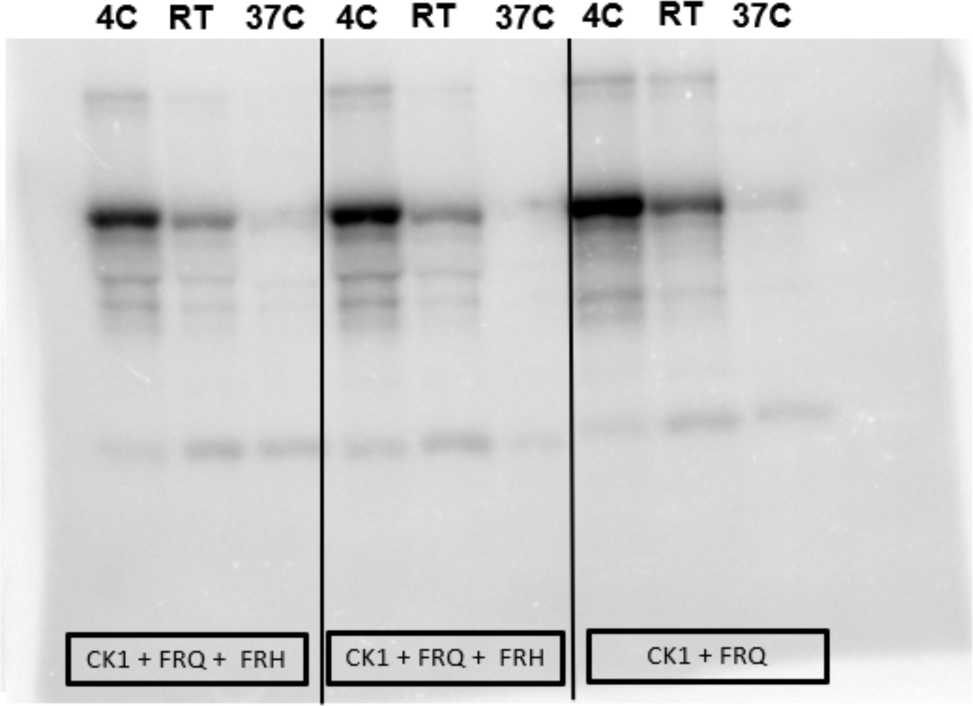
Full radiograph of np-FRQ phosphorylation by CK1 and γ-^32^P-ATP from Figure 7.

**Movie S1:**
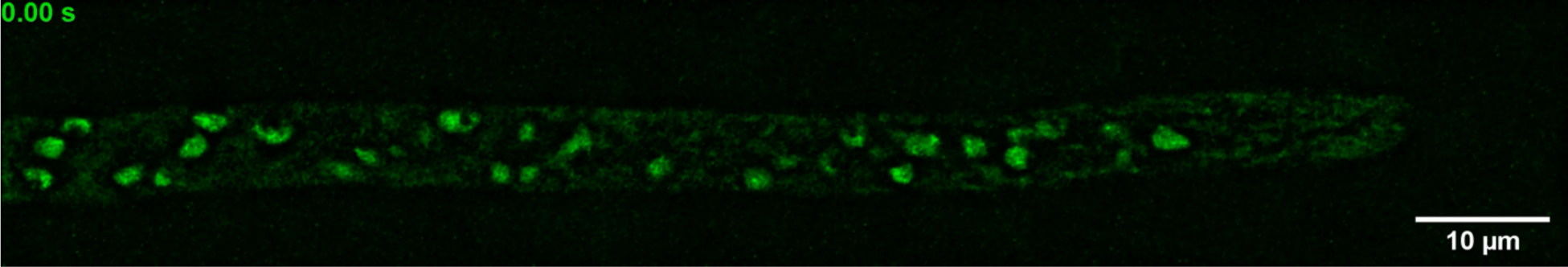
Visualization of FRQ [mNeonGreen] in living Neurospora, displaying heterogeneous, punctate intranuclear localization. Images of a single focal plane were acquired in a time series with 300 ms between exposures.

